# Developmental Dieldrin Exposure Alters DNA Methylation at Genes Related to Dopaminergic Neuron Development and Parkinson’s Disease in Mouse Midbrain

**DOI:** 10.1101/494237

**Authors:** Joseph Kochmanski, Sarah E. VanOeveren, Alison I. Bernstein

**Affiliations:** Department of Translational Science & Molecular Medicine, College of Human Medicine, Michigan State University

**Keywords:** Dieldrin, DNA Methylation, Epigenetics, Parkinson’s Disease, RNA Sequencing

## Abstract

Human and animal studies have shown that exposure to the organochlorine pesticide dieldrin is associated with increased risk of Parkinson’s disease (PD). Despite previous work showing a link between developmental dieldrin exposure and increased neuronal susceptibility to MPTP toxicity in male C57BL/6 mice, the mechanism mediating this effect has not been identified. Here, we tested the hypothesis that developmental exposure to dieldrin increases neuronal susceptibility via genome-wide changes in DNA methylation. Starting at 8 weeks of age and prior to mating, female C57BL/6 mice were exposed to 0.3 mg/kg dieldrin by feeding (every 3 days) throughout breeding, gestation, and lactation. At 12 weeks of age, pups were sacrificed and midbrains were dissected. DNA was isolated and dieldrin-related changes in DNA methylation were assessed via reduced representation bisulfite sequencing (RRBS). We identified significant, sex-specific differentially methylated CpGs (DMCs) and regions (DMRs) by developmental dieldrin exposure (FDR<0.05), including DMCs at the *Nr4a2* and *Lmx1b* genes, which are involved in dopaminergic neuron development and maintenance. Developmental dieldrin exposure had distinct effects on the male and female epigenome. Furthermore, a separate set of changes in DNA methylation was identified after adult exposure to dieldrin, suggesting that adult and developmental dieldrin toxicity may not act through a shared epigenetic mechanism. Together, our data suggest that developmental dieldrin exposure establishes sex-specific poised epigenetic states early in life. These poised epigenomes may mediate sensitivity to additional environmental stimuli and contribute to the development of late-life neurodegenerative disease, including PD.

## 1. Introduction

Parkinson’s disease (PD), the most common neurodegenerative movement disorder, is characterized by formation of α-synuclein (α-syn)-containing Lewy bodies and progressive degeneration of dopaminergic neurons of the nigrostriatal pathway (Fahn, 2003). Previous research has estimated that only 5-10% of PD cases are familial (caused by monogenic mutations), whereas the remaining 90-95% of cases are sporadic (Lill, 2016; Trinh and Farrer, 2013). While the etiology of sporadic PD remains unclear, studies suggest it may involve an interaction between genetics and environment (Lill, 2016; Cannon and Greenamyre, 2013; Fleming, 2017). Supporting this idea, both human and animal studies have shown that exposure to certain classes of environmental toxicants, including specific types of industrial toxicants and pesticides, is associated with increased risk of PD (Freire and Koifman, 2012; Caudle *et al*., 2012; Goldman *et al*., 2017; Moretto and Colosio, 2011; Cicchetti *et al*., 2009).

Dieldrin is a highly toxic organochlorine pesticide that was phased out of commercial use in the 1970s, but has persisted in the environment due to its high stability and lipophilicity (CDC, 2016). Given these properties, dieldrin accumulates in lipid-rich tissues like the brain, and has been classified as a persistent organic pollutant (Corrigan *et al*., 2000; Jorgenson, 2001; Kanthasamy *et al*., 2005). Previous epidemiology studies have shown a positive association between dieldrin exposure and PD risk (Kanthasamy *et al*., 2005; Moretto and Colosio, 2011), and mechanistic animal research has shown that adult and developmental dieldrin exposures are associated with oxidative stress, disrupted expression of PD-related proteins, and increased susceptibility of dopaminergic neurons to toxicants that target the dopaminergic system (Richardson *et al*., 2006; Hatcher *et al*., 2007). In particular, male and female mice developmentally exposed to dieldrin showed increased dopamine transporter (DAT) and vesicular monoamine transporter 2 (VMAT2) protein levels, as well as increased *Nr4a2* expression. In male mice only, developmental exposure to dieldrin led to an increased DAT:VMAT2 ratio and exacerbated MPTP neurotoxicity. However, the biological mechanism mediating these long-lasting effects of developmental dieldrin exposure on the dopaminergic system has not been identified.

The epigenome is recognized as a potential mediator of the relationship between developmental exposures and adult disease. Epigenetic marks are sensitive to the environment, established during cellular differentiation, and regulate gene expression throughout the lifespan (Allis and Jenuwein, 2016; Faulk and Dolinoy, 2011). Thus, it is possible that developmental dieldrin exposure induces fixed changes in the epigenome, creating a poised epigenetic state in which the developmental exposure has programmed a modified response to later-life challenges.

Based on the documented link between developmental dieldrin and adult susceptibility to neurodegeneration, we hypothesized that developmental dieldrin exposure increases neuronal susceptibility to toxicants that target the dopaminergic system via genome-wide changes in DNA methylation. To test this hypothesis, we characterized genome-wide DNA methylation in animals developmentally exposed to dieldrin.

## 2. Materials and Methods

### 2.1 Animals

Male and female C57BL/6 mice were purchased from Jackson Laboratory (Bar Harbor, ME, USA). Mice were maintained on a 12:12 reverse light/dark cycle. Food and water were available *ad libitum*. Mice were maintained on standard LabDiet 5021 chow (LabDiet, St. Louis, MO). F0 females were individually housed during dieldrin dosing, except during the mating phase. F1 pups were group housed by sex; no more than four animals were housed per cage. All procedures were conducted in accordance with the National Institutes of Health Guide for Care and Use of Laboratory Animals and approved by the Institutional Animal Care and Use Committee at Michigan State University.

### 2.2 Dieldrin exposure paradigms

For both adult and developmental exposure, mice were administered 0.3 mg/kg dieldrin dissolved in corn oil vehicle and mixed with peanut butter pellets every 3 days (Gonzales *et al*., 2014). Control mice received an equivalent amount of corn oil vehicle in peanut butter. This dose was used based on previous results showing low toxicity, but clear developmental effects (Richardson *et al*., 2006). In both exposures, animals were fed using a mixture of corn oil and peanut butter, while vehicle control consisted of peanut butter pellets with only corn oil added.

In the adult exposure study, 8-week-old C57BL/6 male animals were treated for 30 days. After the 30-day exposure period, mice were sacrificed and midbrain samples were collected.

In the developmental exposure study, adult C57BL/6 (8-week-old) female animals were treated throughout breeding, gestation, and lactation (**Figure 1A**). Four weeks into female exposure, C57BL/6 males were introduced for breeding. Offspring were weaned at 3 weeks of age, at which point they were separated by litter and by sex. At 12 weeks of age, male and female offspring from independent litters (n=12) were sacrificed and midbrain samples were collected. This time point was chosen based on previous results demonstrating increased neuronal susceptibility to MPTP at 12 weeks of age (Richardson *et al*., 2006).

**Figure 1.**
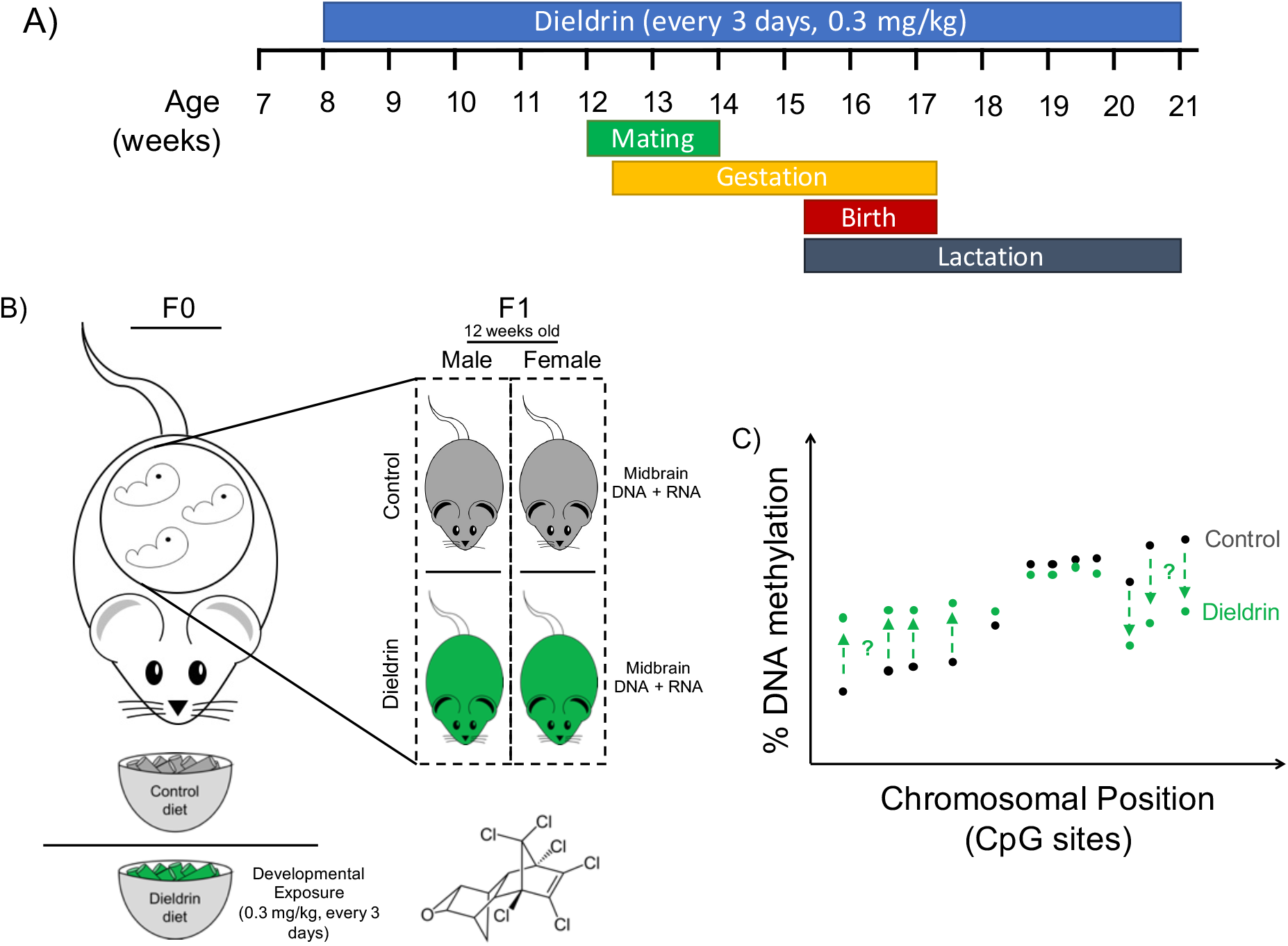
Developmental exposure paradigm and timeline. A) Adult C57BL/6 (8-week-old) female animals were treated with dieldrin (0.3 mg/kg, every 3 days) or vehicle control during the entire developmental period. Dietary dieldrin was fed orally using a mixture of corn oil and peanut butter throughout breeding, gestation, and lactation. Vehicle control mice were fed peanut butter with corn oil only. Four weeks into female exposure, males were introduced for breeding. Offspring were weaned at 3 weeks of age, at which point they were separated by litter and by sex. After developmental exposure, male and female offspring (F1) were raised on control diet and followed to adulthood. B) At 12 weeks of age, F1 offspring were sacrificed, bilateral midbrain samples were dissected, and DNA and RNA were isolated from paired littermates. The transcriptome was assessed from RNA samples using RNA-sequencing technology and genome-wide DNA methylation was assessed by reduced representation bisulfite sequencing. C) Genome-wide DNA methylation was measured from isolated DNA to identify specific cytosine-phospho-guanine (CpG) sites that showed differential DNA methylation by developmental dieldrin exposure.

### 2.3 DNA and RNA isolations

DNA isolation was performed using the phenol:chloroform:isoamyl alcohol method. Briefly, 20 μL proteinase K (Qiagen, Hilden, Germany) and 180 μL ATL lysis buffer (Qiagen, Hilden, Germany) were added to each midbrain tissue sample in a 1.5 mL microcentrifuge tube. A Kimble Kontes pellet pestle (Kimble Chase, Rockwood, TN) was then used to grind the tissue. After vortexing, samples were allowed to digest for 3 hours in a ThermoMixer set at 55°C, 300 rpm. Lysed samples were then centrifuged for 1 min at 12,000 x g and the supernatant was transferred to a new 1.5 mL tube. To ensure RNA-free genomic DNA isolation, 4 μL RNase A (100 mg/mL) (Qiagen, Hilden, Germany) was added to each supernatant; samples were then mixed by vortexing and incubated at room temperature for two minutes. An equal volume of phenol:chloroform:isoamyl alcohol (ThermoFisher, Waltham, MA) was then added to each supernatant. Samples were mixed by inverting for approximately 20 seconds, and then centrifuged at room temperature for 5 minutes at 16,000 x g. The top aqueous phase (containing DNA) was then transferred to a new 1.5 mL microcentrifuge tube. To precipitate DNA, 7.5 M ammonium acetate (Sigma-Aldrich, St. Louis, MO) and 100% ethanol (Decon Labs Inc., King of Prussia, PA) were added at 0.5x and 2.5x the sample volume, respectively. Tubes were then placed in −80°C freezer for 2 hours to precipitate DNA. After thawing samples, DNA was pelleted by centrifugation at 4°C for 30 minutes at 16,000 x g. After removing supernatant, the DNA pellet was washed by adding 150 μL 70% ethanol, and then centrifuging at 4°C for two minutes at 16,000 x g. After repeating this wash step once more, dry pellets were centrifuged again at 4°C for 1 minute at 16,000 x g. Excess 70% ethanol was removed, and tubes were inverted on the benchtop for 5 minutes or until pellets were dry. DNA pellet was then resuspended in 50 μL 10 mM UltraPure Tris pH 8.0 (ThermoFisher, Waltham, MA). DNA yield was determined using a Qubit 3 fluorometer (ThermoFisher, Waltham, MA), and DNA purity was assessed using a Take3 plate micro-volume plate on a BioTek Synergy H1 (BioTek Instruments, Inc., Winooski, VT). Isolated DNA was stored at −80°C.

RNA isolation was performed using a modified RNeasy Lipid Tissue Mini Kit (Qiagen, Hilden, Germany). Several changes were made to the standard RNeasy Lipid Tissue Mini kit to improve RNA yield from midbrain samples. First, tissue was homogenized in 200 μL cold Qiazol lysis reagent using a Kimble Kontes pellet pestle (Kimble Chase, Rockwood, TN) for microcentrifuge tubes. Second, after physical homogenization, an additional 800 μL of Qiazol lysis reagent was added to each tissue sample. Third, after initial centrifugation, supernatant was transferred to an Invitrogen Phasemaker tube (ThermoFisher, Waltham, MA) to facilitate transfer of the RNA-containing aqueous layer. As a result of using the Phasemaker tubes, the second centrifugation step was reduced from 15 minutes to 5 minutes. In addition to these changes, the optional DNase digestion step (RNeasy Lipid Tissue Appendix C) was included to improve purity of isolated RNA. RNA was eluted in 50 μL RNase-free water, and RNA yield and purity were both assessed using the Agilent RNA 6000 Pico Reagents with the Agilent 2100 Bioanalyzer System (Agilent Technologies, Santa Clara, CA). Isolated RNA was stored at −80°C.

### 2.4 Reduced Representation Bisulfite Sequencing

Genome-wide DNA methylation levels were measured using reduced representation bisulfite sequencing (RRBS) (Gu *et al*., 2010). RRBS is a genome-wide sequencing method that utilizes a restriction enzyme (*MspI*) to specifically cuts at CCGG nucleotide sequences, thereby enriching for CpG-dense regions. RRBS library preparation was performed at the University of Michigan Epigenomics Core. Briefly, 200 ng of genomic DNA was digested for 16-18 hours using MspI, a restriction enzyme that preferentially cuts DNA at CCGG nucleotide sequences. Restriction enzyme-digested DNA was purified using phenol:chloroform extraction and ethanol precipitation in the presence of glycogen and sodium acetate. Precipitated DNA was resuspended in 10 mM Tris-Cl pH 8.5 and processed for end-repair, A-tailing, and ligation of adapters with methylated cytosines. The ligated DNA was further cleaned according to the NuGEN TrueMethyl oxBS module (NuGEN, Redwood City, CA; Cat # 0414-32) instructions with 80% acetonitrile and magnetic beads. Ligated DNA was then eluted in UltraPure water and split into two tubes, one processed for bisulfite conversion only and the other processed for oxidative-bisulfite treatment, according to the TrueMethyl kit’s instructions. After bisulfite conversion and cleanup, the samples were amplified using the Roche High Fidelity FastStart system ((Sigma-Aldrich, St. Louis, MO) and TruSeq PCR primers (Illumina Inc., San Diego, CA), according to the following protocol: 94°C for 5 minutes; 18 cycles of 94°C for 20 seconds, 65°C for 30 seconds, and 72°C for 1 min; 72°C for 3 minutes. Final library cleanup was done with 1:1 volume of Agencourt AMPure XP SPRI beads (Beckman Coulter, Brea, CA). Each library was eluted into 20 μl of 10 mM Tris-HCl pH 8.5. Libraries were quantified using the Qubit dsDNA High Sensitivity kit (ThermoFisher, Waltham, MA), and library quality was assessed with the Agilent High Sensitivity D1000 ScreenTape assay (Agilent, Santa Clara, CA). Sequencing of RRBS libraries was performed at the Van Andel Genomics Core. To minimize batch effects, individually indexed libraries were randomized and pooled into two groups. Paired-end, 75 bp sequencing was performed on an Illumina NovaSeq6000 sequencer using 200 bp S1 sequencing kits (Illumina Inc., San Diego, CA). RRBS data had genomic coverage of approximately 3,000,000 CpG sites for each sample.

### 2.5 RNA-sequencing

Whole-genome RNA transcript expression was measured using RNA sequencing (RNA-seq) (Wang *et al*., 2009). RNA-seq library preparation was performed at the Van Andel Genomics Core. Libraries were prepared from 100 ng of total RNA using the KAPA RNA HyperPrep Kit with RiboseErase (v1.16) (Kapa Biosystems, Wilmington, MA USA). As part of the library prep, RNA was sheared to 300-400 bp. After cDNA synthesis, but prior to PCR amplification, cDNA fragments were ligated to Bioo Scientific NEXTflex Adapters (Bioo Scientific, Austin, TX, USA). Library quality and quantity was assessed using a combination of Agilent DNA High Sensitivity chip (Agilent Technologies, Inc.), QuantiFluor^®^ dsDNA System (Promega Corp., Madison, WI, USA), and Kapa Illumina Library Quantification qPCR assays (Kapa Biosystems). Individually indexed libraries were randomized and pooled to minimize batch effects. Paired-end, 75 bp sequencing was performed on an Illumina NextSeq 500 sequencer using a 150 bp HO sequencing kit (v2) (Illumina Inc., San Diego, CA, USA). All libraries were run across 3 flowcells to achieve a minimum read depth of 60M read pairs per library.

### 2.6 Sequencing QC and adapter trimming

Bioinformatics pipelines are outlined in **Figure 2**. Linux command line tools and the open-source statistical software R (version 3.5.1) were used for all analyses. For all sequencing data, the *FastQC* tool (version 0.11.5) was used for data quality control, and the *trim_galore* tool (version 0.4.5) was used for adapter trimming (Krueger, 2017; Andrews, 2016). During adapter trimming, we used the default minimum quality score and added a stringency value of 6, thereby requiring a minimum overlap of 6 bp. For the RRBS data, reads were trimmed in “RRBS mode” using the --rrbs parameter; this parameter was not included for RNA-seq data trimming.

**Figure 2.**
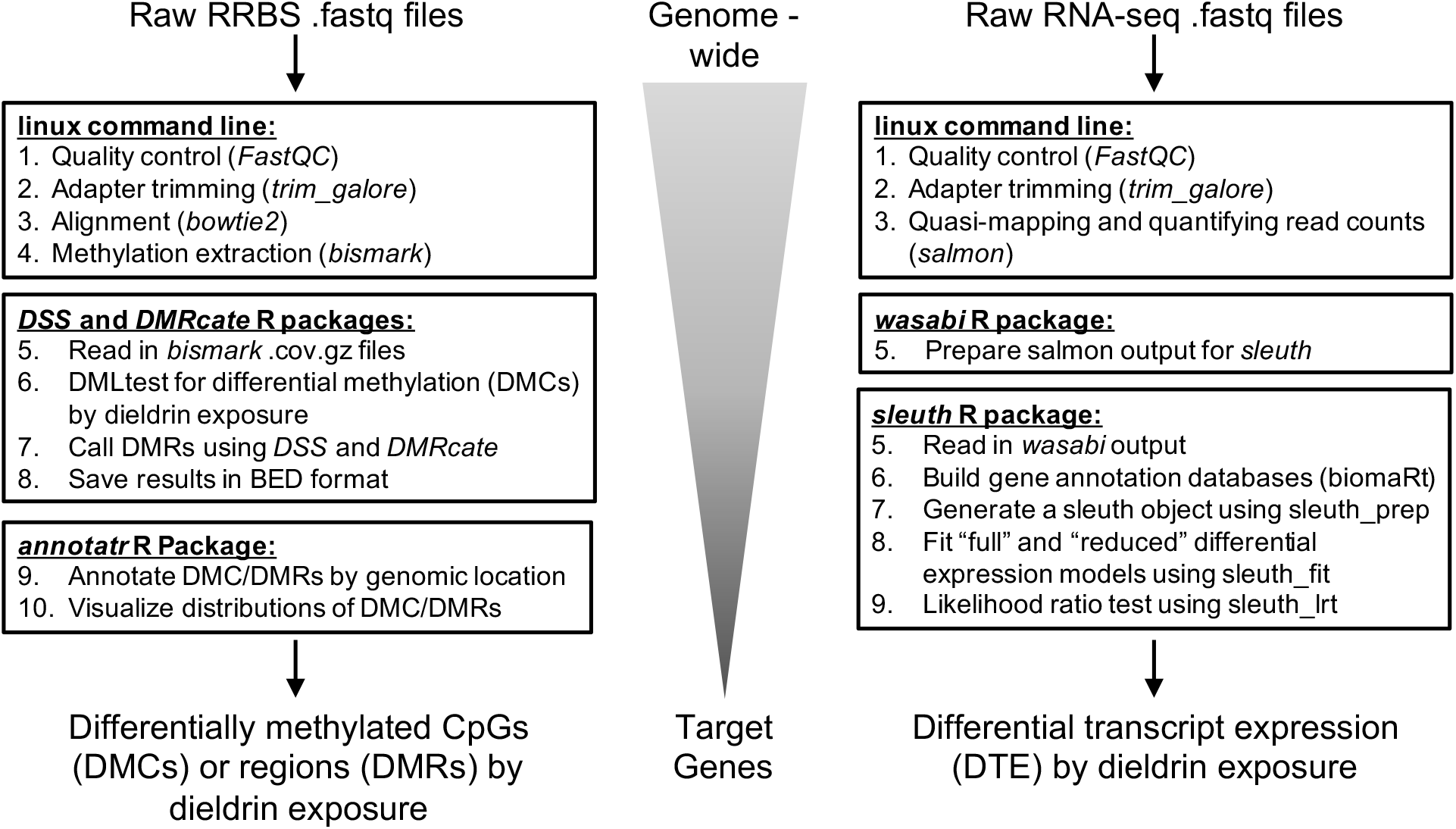
Reduced representation bisulfite sequencing (RRBS) and RNA-sequencing (RNA-seq) bioinformatics pipelines. RRBS and RNA-seq data were separately processed and analyzed using the indicated bioinformatics pipelines. The goal of the pipelines was to identify genome-wide changes in DNA methylation (RRBS) or transcript expression (RNA-seq) by developmental dieldrin exposure, and then generate lists of target genes. Linux command line tools and the open-source statistical software R (version 3.5.1) were used for all analyses.

### 2.7 Differential methylation analysis

Trimmed RRBS reads were aligned to the mm10 genome using *bismark* (version 0.19.1) and *bowtie2* (version 2.3.2) with default parameters (Langmead, 2017; Krueger, 2018). Methylation data was extracted from the aligned reads in *bismark* using a minimum threshold of 5 reads to include a CpG site in analysis.

The *DSS* (version 2.30.0) and *DMRcate* (version 1.18.0) R packages were used in parallel to test RRBS data for differential methylation by dieldrin exposure (Feng *et al*., 2014; Peters *et al*., 2015). Given that dieldrin exposure has shown sex-specific effects on the dopaminergic phenotype (Richardson *et al*., 2006), all differential methylation models were stratified by sex. To test for differentially methylated CpGs (DMCs) by dieldrin exposure, we used the DMLtest function in *DSS* to perform two-group Wald tests. For DMLtest modeling, the equal dispersion and smoothing parameters were set to false. DMC significance was determined using a false discovery rate (FDR) cutoff <0.05. To test for differentially methylated regions (DMRs), we combined outputs from the callDMR function in *DSS* and the dmrcate function in *DMRcate*. For callDMR modeling, the p-value threshold was set to 0.05, minimum length was set to 20 base pairs, and minimum CpGs was set to 3. Meanwhile, for dmrcate modeling, the lambda value was set to 500, the C value was set to 4, and minimum CpGs was set to 3. Within the dmrcate function, groups of DSS-derived FDRs for individual CpG sites are Stouffer transformed; here, DMR significance for the dmrcate output was set to Stouffer value < 0.10.

After differential methylation testing, the *annotatr* R package (version 1.8.0) was used to annotate identified DMCs and DMRs to the reference mm10 genome (Cavalcante and Sartor, 2017). Within *annotatr*, the annotate_regions function was used to generate CpG category, gene body, and regulatory feature annotations. To provide a more complete annotation, custom miRNA (sourced from: http://www.mirbase.org/ftp.shtml) and ENCODE predicted mouse midbrain enhancer (sourced from: https://www.encodeproject.org/annotations/ENCSR114ZIJ/) databases were added to the annotation cache in *annotatr*. Distributions of DMCs by genomic annotation were plotted using the plot_annotation function in *annotatr* (**Figure S1**). Overlap between male- and female-specific DMCs and DMRs was tested using the bedtools (version 2.27.1) intersect function (Quinlan and Kindlon, 2017).

### 2.8 Differential expression analysis

Following adapter trimming, read counts were quantified and quasi-mapped to a self-generated mm10 index using the *salmon* tool (version 0.11.3) (Patro *et al*., 2017). During read quantification in *salmon*, the --numBootstraps parameter was used to generate 100 bootstraps for each sample. Quantified RNA-seq reads were then processed for downstream analyses using the prepare_fish_for_sleuth function in the *wasabi* R package (version 0.3) (Patro, 2018).

The *sleuth* R package (version 0.30.0) was used to test wasabi-processed RNA-seq data for differential transcript expression (DTE) by dieldrin exposure (Pimentel *et al*., 2017). Matching the DNA methylation analysis pipeline, all DTE models were stratified by sex. Prior to analysis, the Ensembl mouse transcript annotation database was read into R using the biomaRt package. Additionally, to limit analyses to high-quality data, only transcripts with greater than 10 estimated counts in at least 50% of the samples were included in downstream steps. After loading annotations and filtering by count, DTE models were generated in the *sleuth* package using a combination of the sleuth_prep, sleuth_fit, and sleuth_lrt functions. The sleuth_prep function initialized the sleuth object and read in the target mapping and bootstrap information from the provided annotation database and wasabi-processed data, respectively. The sleuth_fit function was then used to produce two smoothed linear models – “full” and “reduced.” In *sleuth*, the “full” model was fit using a smoothed linear model for experimental treatment (control vs. dieldrin), whereas the “reduced” model was fit assuming equal abundances by treatment. Finally, the sleuth_lrt function uses a likelihood ratio test (LRT) to identify transcripts with a significantly improved fit in the “full” model. DTE significance for the *sleuth* LRT was set to FDR < 0.05.

### 2.9 Data visualization

RRBS data were visualized using the *RnBeads* R package (version 2.0.0) (Assenov *et al*., 2014). First, the rnb.execute.import function was used to import the processed RRBS data. Next, the rnb.sample.groups function was used to order samples by treatment group. Finally, the rnb.plot.locus.profile function was used to plot DNA methylation values for specific regions of the genome (i.e. identified DMCs or DMRs). For RNA-seq data, estimated counts extracted from *sleuth* were visualized using the *ggplot2* R package (version 3.1.0).

### 2.10 Pathway and Network Analysis

Gene ontology (GO) term enrichment testing and pathway analysis was performed on genes annotated to male and female DMCs using the ClueGO application in Cytoscape (version 3.6.1) (Bindea *et al*., 2009; Smoot *et al*., 2011). For ClueGO testing, hypo- and hypermethylated DMCs were input as separate marker lists, “groups” was selected as the visual style, and the biological processes GO term was included for enrichment testing. Network specificity was set half-way between “Medium” and “Detailed”, such that the GO Tree Interval minimum was equal to 6 and the maximum was equal to 12. Only terms with at least 3 genes and a Bonferroni-corrected p-value < 0.1 were included in pathway visualizations. The connectivity score (Kappa) was set at 0.5, and default GO Term Grouping settings were used in all analyses. In addition to ClueGO analyses, gene-gene interaction network analysis was performed on genes annotated to male and female DMCs using STRING (version 10.5) (Szklarczyk *et al*., 2017). STRING network analysis was performed using default parameters, including a minimum required interaction score = 0.4 and all interaction sources activated.

## 3. Results

### 3.1 Differential methylation by developmental dieldrin exposure

The *DSS* R package was used to identify differentially methylated cytosines (DMCs) from the RRBS data using a Bayesian hierarchical model based on the beta-binomial distribution. Due to the sex-specific phenotypic effects of dieldrin on neuronal susceptibility, we stratified differential methylation models by sex. Two-group Wald test models were used to test for differential DNA methylation by developmental dieldrin exposure (**Figure 3A**). In the female mice, we identified 478 DMCs (**Table S1**). Comparing the directionality of the female DMCs, more DMCs were hypomethylated (n=321; 67.2%) than hypermethylated (n=157; 32.8%), suggesting a general pattern of decreased methylation with dieldrin exposure in female mice. Meanwhile, in the male mice, we identified 115 DMCs (**Table S2**). Unlike the female data, more the male DMCs were hypermethylated (n=74; 64.3%) than hypomethylated (n=41; 35.7%), indicating a pattern of hypermethylation in the male mice. Male and female DMCs showed no overlap between the two sexes (**Figure 3C**).

**Figure 3.**
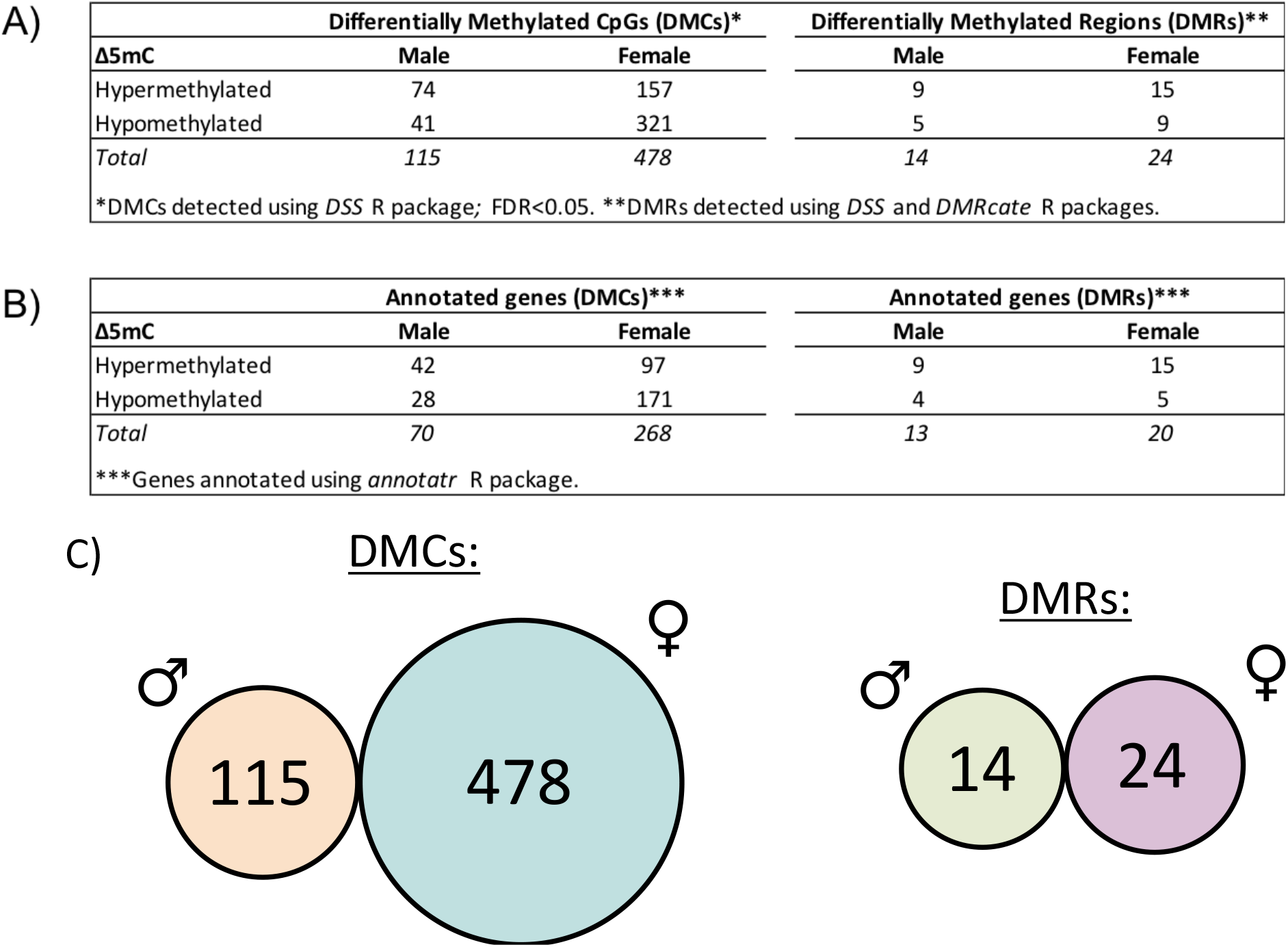
Dieldrin-related differential methylation identified from RRBS data. Using the *DSS* R package, two-group Wald test models were used to test for differential DNA methylation by developmental dieldrin exposure in RRBS data from mouse midbrain samples collected at 12 weeks of age. Due to the sex-specific phenotypic effects of dieldrin on neuronal susceptibility, we stratified differential methylation models by sex. A) In female mice, we identified 478 differentially methylated CpGs (DMCs) by dieldrin exposure. More DMCs were hypomethylated (n=321; 67.2%) than hypermethylated (n=157; 32.8%), indicating a general pattern of decreased methylation with dieldrin exposure in female mice. Meanwhile, in the male mice, we identified 115 DMCs by dieldrin exposure. Unlike the female data, more of the male DMCs were hypermethylated (n=74; 64.3%) than hypomethylated (n=41; 35.7%), indicating a general pattern of hypermethylation in the male mice. B) DMCs and DMRs were annotated to genes using the *annotatr* R package. Not all identified DMCs and DMRs annotated to known gene regions. C) The bedtools intersect function was used to test for overlap between male and female DMCs/DMRs by chromosomal location. Male and female DMCs/DMRs showed no overlap between the two sexes.

The *DSS* and *DMRcate* R packages were used to test for larger regions of differential methylation by dieldrin exposure (**Figure 3A**). In the female mice, we found 24 differentially methylated regions (DMRs) by dieldrin exposure (**Table S3**). Meanwhile, in the male mice, we identified 14 DMRs by dieldrin exposure (**Table S4**). In contrast to DMCs, for both the male and female mice, there was a slight skew toward hypermethylated DMRs (female n=15; male n=9) compared to hypomethylated DMRs (female n=9; male n=5). Consistent with our observations regarding the DMCs, the male and female DMRs showed no overlap between the two sexes (**Figure 3C**), showing sex-specific patterns of differential methylation on the regional scale.

The *annotatr* R package was used annotate the detected DMCs and DMRs to known murine genes, regulatory features, enhancers and miRNAs (**Figure 3B**). DMCs and DMRs that annotated to any region or feature of a gene were considered “annotated” to that gene. In the female mice, the identified DMCs and DMRs annotated to 268 and 20 genes, respectively (**Tables S5 and S6**). Meanwhile, in the male mice, the identified DMCs and DMRs annotated to 70 and 13 genes, respectively (**Tables S7 and S8**). Reflecting the lack of overlap between DMCs and DMRs, we also observed no overlap between annotated genes. In addition, we also characterized the functional annotation of the DMCs, annotating to known CpG categories (e.g. CpG island), gene body locations (e.g. exon), and regulatory features (e.g. promoter). Both male and female DMCs showed similar genomic distributions, indicating that while specific patterns of dieldrin-induced differential methylation were sex-specific, the overall categories of features affected by exposure were not necessarily different by sex (**Figure S1**).

### 3.2 ClueGO and STRING pathway/network analyses for developmental exposure model

Given that the DMCs annotated to a large number of genes, ClueGO was used to perform a gene ontology enrichment analysis on the genes with dieldrin-induced differential methylation (**Figure 4A**). Input lists for ClueGO included all genes with annotated DMC or DMRs, regardless of DMC/DMR location. For the male DMCs, we did not identify any significantly enriched GOBP terms after multiple testing correction. In contrast, for the female DMCs, we found a large number of significantly enriched gene ontology-biological process (GOBP) terms, including cranial nerve development and central nervous system neuron development (Bonferroni-corrected p-value <0.05) (**Figure 4B**). Most of the other enriched terms were also related to development, although in different biological systems (**Table S9**). Next, the genes included in the significant GOBP terms (listed in **Figure 4B**) were extracted and placed into the STRING network interaction tool to investigate interactions between these genes (**Figure 4C**). We found that the genes from the significant female GOBP terms clustered together into a network. Of particular interest, two of the differentially methylated genes from the central nervous system neuron development pathway – *Nr4a2* and *Lmx1b* – are known to be critical for dopaminergic neuron development and maintenance.

**Figure 4.**
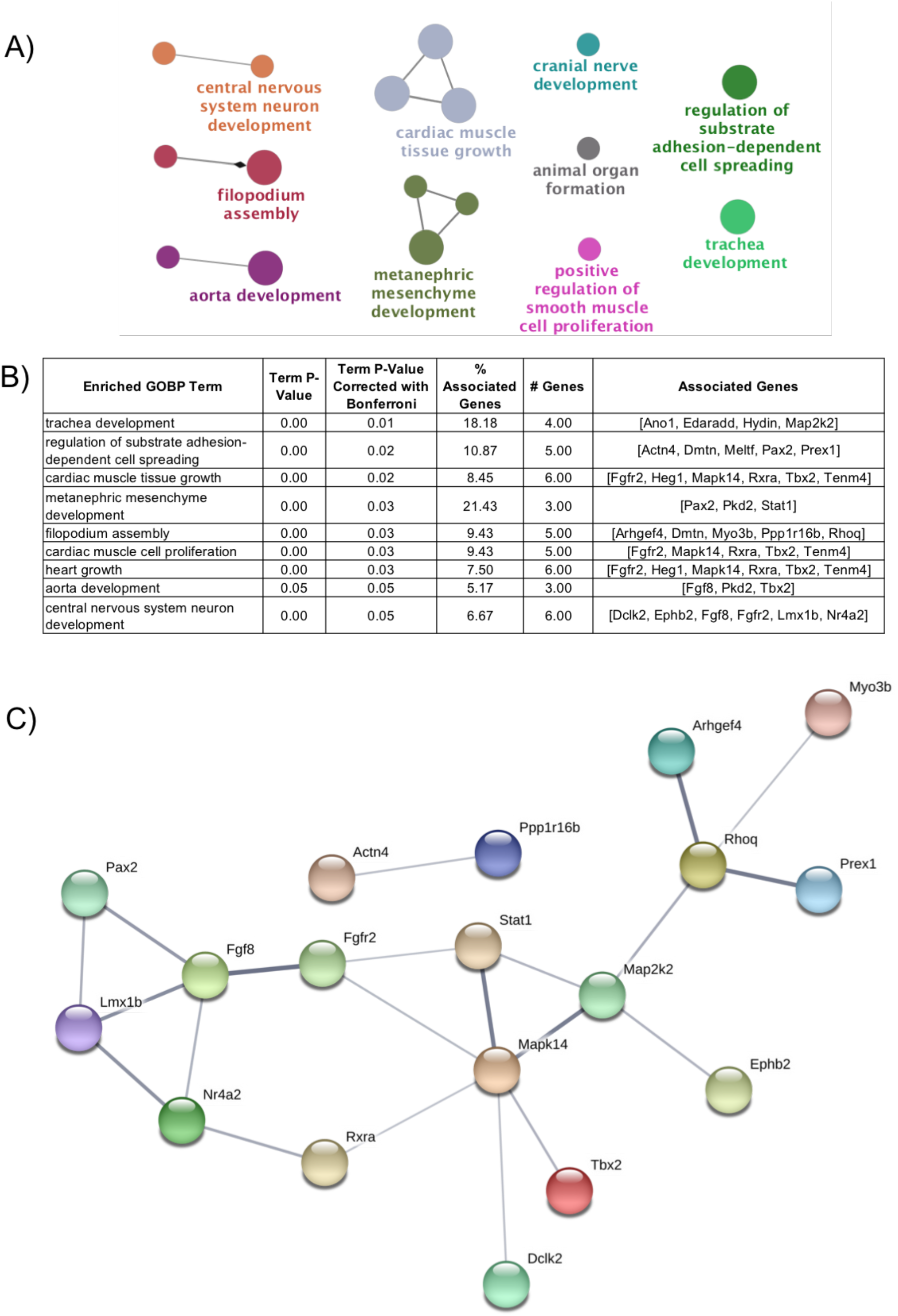
ClueGO pathway analysis, significant terms/genes, and STRING interaction network. A) ClueGO was used to perform a gene ontology enrichment analysis on the genes with dieldrin-induced differential methylation. For the male DMCs, we did not identify any significantly enriched GOBP terms after correction for multiple testing. However, for the female DMCs, we found a large number of significantly enriched gene ontology-biological process (GOBP) terms, including cranial nerve development and central nervous system neuron development (Bonferroni-corrected p-value <0.05). B) Genes included in the significant GOBP terms were extracted. C) Genes from enriched GO terms were placed into the STRING network interaction tool to investigate interaction amongst these genes. We found that the genes from the significant female GOBP terms clustered together into a network. Two of the differentially methylated genes in the network – *Nr4a2* and *Lmx1b* – are known to be involved in dopaminergic neuron development. Six of the genes – *Heg1*, *Ano1*, *Edaradd*, *Hydin*, *Pkd2*, and *Tenm4* – showed no interactivity in the STRING network and were removed from the visualization.

### 3.3 Differential methylation at target genes from developmental exposure model

To follow-up on the pathway and network analyses, the *RnBeads* R package was used to visualize DNA methylation levels at all differentially methylated genes in the central nervous system neuron development pathway. While most of the genes showed minimal widespread differential methylation across visualized gene bodies (data not shown), more prevalent changes in DNA methylation were apparent at the *Nr4a2* and *Lmx1b* genes, which have been implicated in dopaminergic neuron development and potentially Parkinson’s disease (Doucet-Beaupré *et al*., 2016; Luo, 2012; Dong *et al*., 2016). Although only a single DMC was annotated to *Nr4a2*, when the entire gene body was visualized, there was a larger region of differential methylation that became apparent (**Figure 5A**). Similarly, in *Lmx1b*, we detected only a single DMC 1-5 kb upstream of the *Lmx1b* transcription start site using our statistical methods, but when visualized, there was consistent hypomethylation surrounding the detected DMC (**Figure 5B**). Fitting with the sex-specific genome-wide results, dieldrin-induced hypomethylation at *Nr4a2* (gene body) and *Lmx1b* (exon) was only present in the female mice. The inability of our selected analysis methods to detect these regions as DMRs was likely a result of the stringent cutoffs used by the *DSS* and *DMRcate* packages.

**Figure 5.**
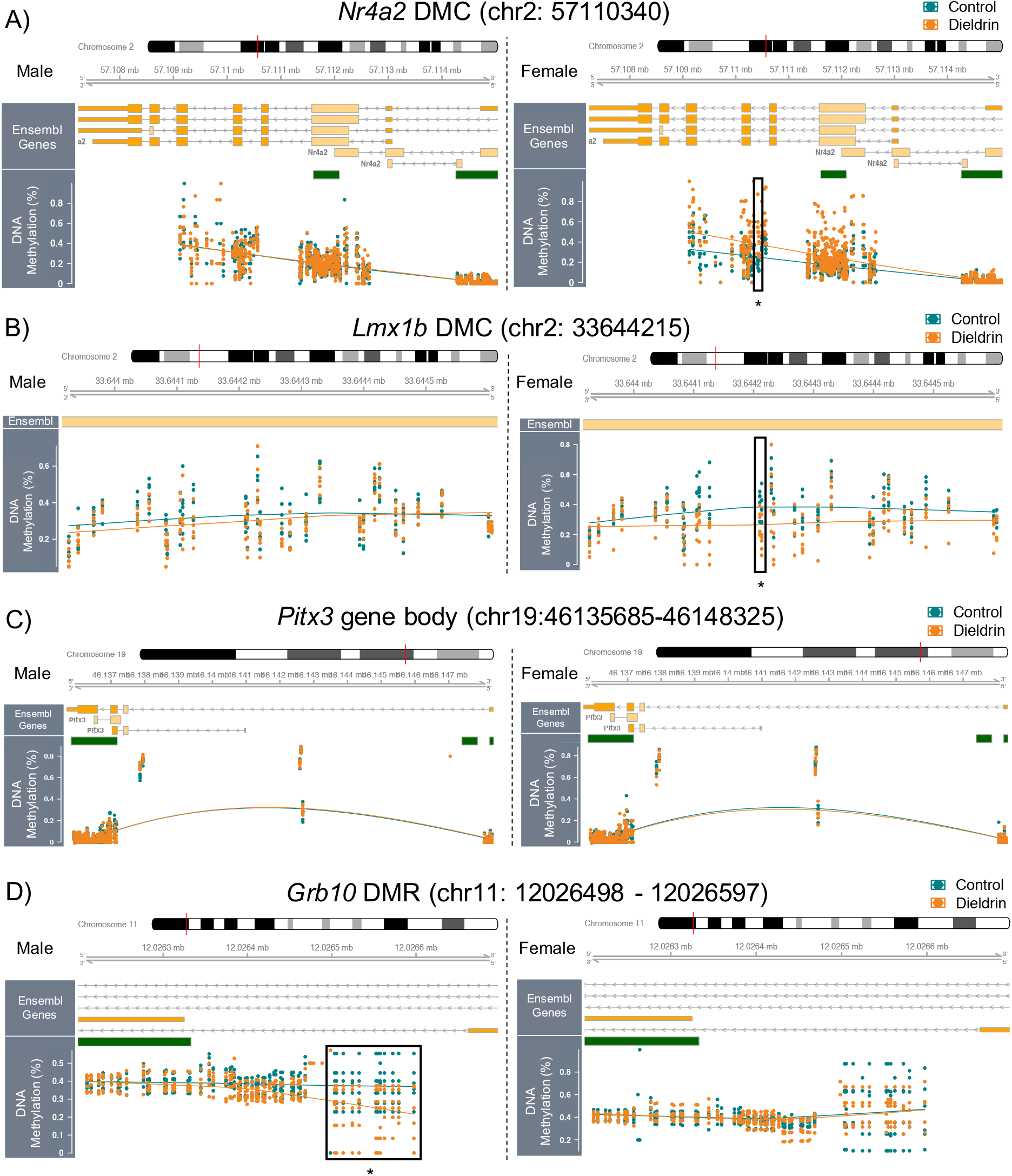
Dieldrin-induced differential methylation at the *Nr4a2, Grb10*, and *Lnvclb* genes. DNA methylation was visualized at the *Nr4a2*, *Grb10*, *Lmx1b*, and *Pitx3* genes using the *RnBeads* and *fíiSeq* R packages. Differentially methylated CpGs (DMCs) and regions (DMRs) were detected using a combination of the *DSS* and *DMRcate* R packages. For all visualizations, the following features are shown from top to bottom: Ensembl gene cartoons, CpG islands (green bars), and DNA methylation levels by individual sample in a dot plot with smoothed mean line. Control and dieldrin group data are shown in green and orange, respectively. As indicated by labels, male data are shown on the left, while female data are shown on the right. Boxes with asterisks indicate areas of differential methylation detected in the *DSS* R package. A) A female-specific intronic DMC (chr2: 57110340, boxed) was identified within *Nr4a2*. The entire *Nr4a2* locus was visualized to show a larger region of apparent, but non-significant, dieldrin-induced hypermethylation surrounding the DMC. B) A female-specific, exonic DMC (boxed) was identified in the *Lmx1b* exon (chr2: 33644215). Additionally, a 700 bp region surrounding the identified *Lmx1b* DMC showed dieldrin-induced hypomethylation in the female mice. C) There was no significant differential methylation at covered CpG sites along the *Pitx3* locus; however, as shown in the plot, coverage across the *Pitx3* gene length was poor. Asterisks below boxes indicate significant differential methylation detected using the *DSS* package (DMC FDR<0.05; DMR p<0.05). D) A male-specific, intronic DMR (boxed) was identified at the imprinted *Grb10* locus (chr11: 12026498 – 12026597).

In addition to the genes that were identified from our pathway and network analyses, we also investigated potential differential methylation at the *Pitx3* gene, a related transcription factor important for dopaminergic neuron development and maintenance than has also been implicated in PD (Li et al. 2009). Unlike *Lmx1b* and *Nr4a2*, however, there were no detected DMCs or DMRs annotated to *Pitx3*, and when visualized, the *Pitx3* gene body showed no apparent differential methylation on a regional scale (**Figure 5C**). However, it should be noted that RRBS data coverage across this gene region was poor apart from the CpG islands, a result that is likely due to the bias of the RRBS method towards CpG-dense regions of the genome.

Given that differentially methylated regions at imprinted genes have shown specific vulnerability to other developmental exposures (Plasschaert and Bartolomei, 2014; Marsit, 2015), we also investigated whether any of our identified dieldrin-related DMCs and DMRs annotated to imprinted loci. One of the identified female DMCs annotated to the promoter of the imprinted *Ascl2* locus, but none of the female DMRs annotated to an imprinted gene. Meanwhile, none of the identified male DMCs annotated to an imprinted gene, but two of the male-specific DMRs annotated to imprinted loci – *Grb10* and *Gnas*. When visualized, the *Gnas* DMR appeared to be a false positive driven by a low number of samples with data coverage (**Figure S2**), but the *Grb10* DMR showed clear hypomethylation with dieldrin exposure in the male midbrain samples (**Figure 5D**). After annotation, we also found that this DMR overlaps a predicted mouse midbrain ENCODE enhancer region.

### 3.4 Whole-genome differential transcript expression for developmental exposure model

Using the *sleuth* R package, a likelihood ratio test was used to test for differential gene expression (DGE) and differential transcript expression (DTE) by dieldrin exposure. After filtering out transcripts with low estimated counts (<10 in more than half of the samples), 56,608 and 55,623 transcripts were included for male and female differential models, respectively. After pooling gene transcripts into a single identifier for DGE models, zero genes showed significant differential expression (FDR<0.10) by dieldrin exposure. Expanding the analysis to the transcript level, there were also zero transcripts with significant differential expression by dieldrin exposure (FDR<0.10) in our genome-wide analyses for male (**Table S10**) and female mice (**Table S11**).

#### 3.4.1 Target gene transcript-level analysis for developmental exposure model

Based on our differential methylation data, pathway analyses, and *a priori* hypotheses generated from previous developmental dieldrin studies, we narrowed the scope of the DTE analysis to the following individual genes: *Nr4a2*, *Lmx1b*, *Grb10*, and *Pitx3*. At the *Nrfa2* gene, one of the protein-coding transcripts showed marginally significant decreased expression in female midbrains by dieldrin exposure (p=0.06) (**Figure 6**). This result was not seen in males (p>0.10), which reflects the observed female-specific differential methylation detected within the *Nr4a2* gene body. These data partially corroborate findings from Richardson et al., who showed nonsignificant increases in *Nr4a2* with 0.3 mg/kg dieldrin in both male and female mice. However, these previous data were generated using primers that measure two protein-coding forms of *Nr4a2* simultaneously (Richardson *et al*., 2006), one of which is presented in **Figure 6**. As such, the female-specific significance presented here may be driven by changes in a specific transcript that could not be detected in previous work. At the *Lmx1b* gene, one protein-coding transcript showed marginally significant decreased expression in male midbrains by dieldrin exposure (p=0.08); this result was not apparent in the female mice (p=0.59) (**Figure 6**). Meanwhile, at the *Grb10* gene, two protein-coding transcripts showed significant and marginally significant decreased expression in male midbrains by dieldrin exposure (p=0.05 and p=0.06, respectively) (**Figure 6**). In contrast, these same two transcripts showed significant dieldrin-related increases in expression in the female midbrains (p<0.05). These results reinforce the idea that dieldrin has sex-specific effects on *Grb10* regulation. Finally, *Pitx3* did not show any differential transcript expression in male or female midbrains (p>0.1) (**Figure 6**), a result that matches the lack of significant differential methylation at this locus. This lack of significant differential expression at the *Pitx3* locus was also consistent with findings from Richardson et al., who showed no change in *Pitx3* expression with 0.3 mg/kg dieldrin in males or females (Richardson *et al*., 2006). These previous data were generated using primers specific to the 1380 bp *Pitx3* protein-coding transcript (Richardson *et al*., 2006), the same transcript presented in **Figure 6**.

**Figure 6.**
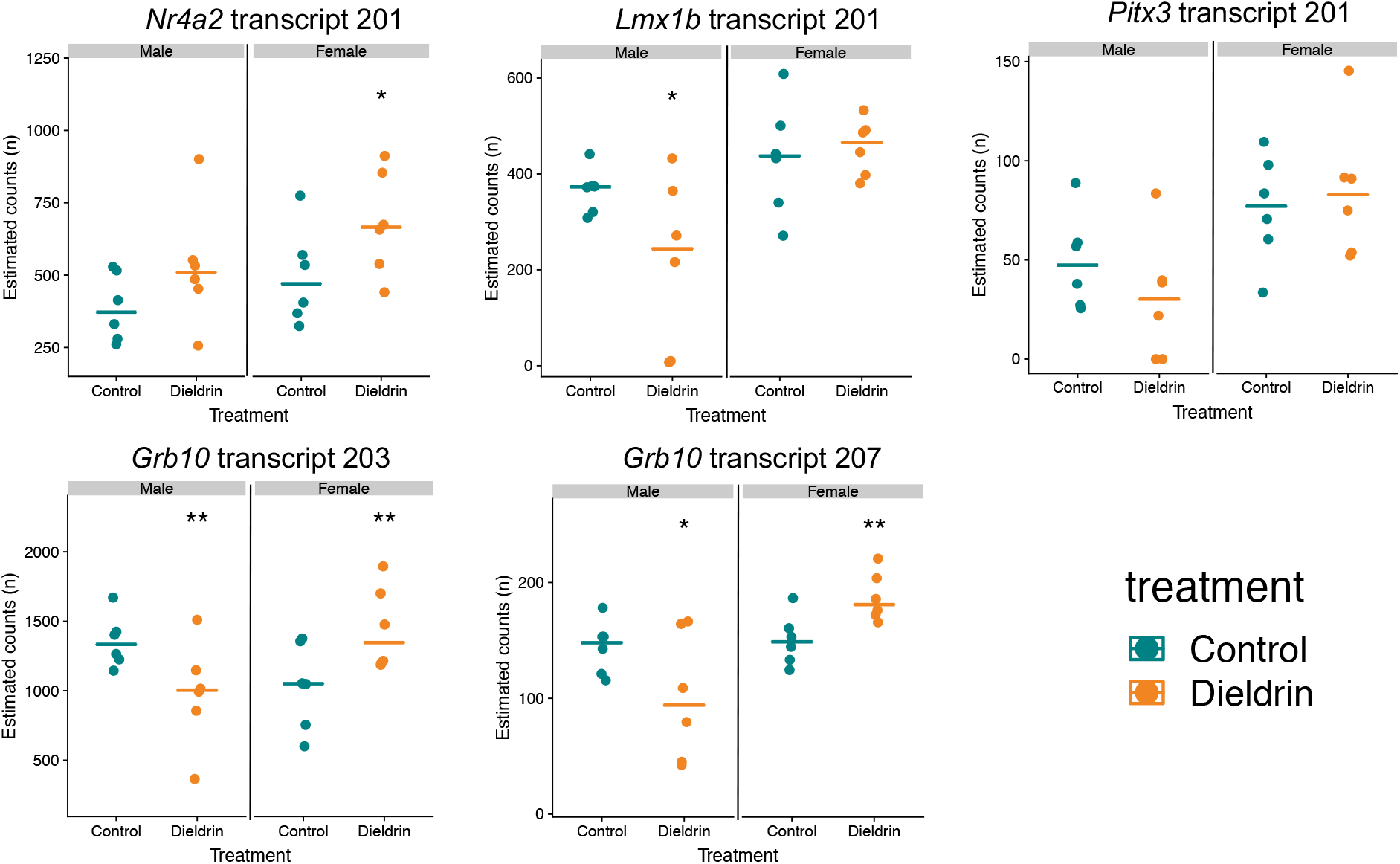
*Nr4a2*, *Grbl0*, *Lnvclb*, and *Pitx3* RNA-seq transcript expression. Dot plots show RNA transcript expression for protein-coding *Nr4a2, Lmxlb*, *Pitx3*, and *Grbl0* transcripts. Control and dieldrin-exposed mice for male and female mice are indicated on the left and right of each plot, respectively. Transcripts were annotated and assigned numbers using the Ensembl mouse transcriptome database (https://useast.ensembl.org/index.htmn. A likelihood ratio test was performed in the *sleuth* R package to determine significance of differential transcript expression. Significance is indicated by asterisks (* p-value ≤ 0.10; ** p-value ≤ 0.05).

### 3.5 Differential methylation in the adult dieldrin exposure model

Using the *DSS* R package, two-group Wald test models were used to test for differential DNA methylation by adult dieldrin exposure in RRBS data generated from male midbrain samples collected at 12 weeks of age. In the adult exposure study, we identified 99 differentially methylated CpGs (**Table S12**), of which 54 were hypermethylated and 45 were hypomethylated (**Figure S3**). In addition to the DMCs, 9 DMRs were detected by adult dieldrin exposure (**Table S13**). The identified DMCs and DMRs annotated to 79 and 4 unique genes, respectively (**Tables S14 and S15**). None of the adult male exposure DMCs or DMRs overlapped with the developmental male exposure model results (**Figure S3**).

Genes with annotated DMCs in the adult exposure model showed little enrichment for GOBP terms after correction for multiple testing. Specifically, only one GO term was enriched in the adult male DMCs – growth hormone secretion. These limited results make it difficult to interpret the biological significance of the adult exposure data. However, the available results do suggest that there are subtle epigenetic effects of dieldrin exposure in adult male mice that are distinct from the effects observed in the developmental exposure model.

## 4. Discussion

### 4.4 Sex-specific epigenome-wide differential methylation by developmental dieldrin exposure

Previous animal work has shown that developmental dieldrin exposure has specific phenotypic effects in mice, including disrupted expression of PD-related proteins, increased dopamine turnover, and increased susceptibility of dopaminergic neurons to the toxicant MPTP (Richardson *et al*., 2006). Here, we investigated whether these phenotypic effects of dieldrin exposure are driven by programmed epigenetic changes. Contrary to expectations, while male mice had previously shown a sex-specific sensitivity to developmental dieldrin exposure (Richardson *et al*., 2006), our identified male DMCs showed no enrichment for pathways related to dopaminergic neuron development. These data suggest that changes in DNA methylation may not be driving the neuronal susceptibility induced by developmental dieldrin exposure in male mice. However, it is also possible that the effects of dieldrin exposure on the male epigenome were simply not captured by the RRBS method, which is biased toward CpG-dense regions. Future studies should utilize whole-genome bisulfite sequencing (WGBS) to more comprehensively answer this question.

Whereas female mice had previously shown a diminished DAT:VMAT2 response to developmental dieldrin exposure compared to males (Richardson *et al*., 2006), we found that female mice had an greater epigenetic response to developmental dieldrin exposure than their male counterparts (**Figure 3**). Furthermore, the female DMCs annotated to genes enriched for a number of gene ontology pathways, including central nervous system development and other developmental processes. These results indicate that the female mouse epigenome is more dynamic in response to dieldrin exposure, a result that is counterintuitive given previous data showing male-specific increases in susceptibility to MPTP after dieldrin exposure. However, it is possible that the pathways affected in female mice are not involved in neurotoxicity, but rather reflect a programmed epigenetic state that maintains dopaminergic function. Future work should examine whether dieldrin-exposed female mice have an improved phenotypic response to alternate Parkinson’s disease models, especially models in which the mechanism of toxicity is not dependent upon alterations in DAT and VMAT2.

Despite a lack of clarity regarding how this dynamic epigenetic regulation in females may alter phenotype, our data add to a growing literature demonstrating the sex-specific nature of epigenetic responses to developmental exposures (Heijmans *et al*., 2008; Leung *et al*., 2018; Kippler *et al*., 2013; Kundakovic *et al*., 2013). Given that the male and female epigenomes can respond so differently to the environment, it is crucial that developmental exposure studies investigating epigenetic effects include both sexes.

### 4.2 Dieldrin-related differential methylation at genes related to dopaminergic neuron development

Using ClueGO pathway analysis, we found that the female DMCs were enriched at genes related to developmental processes, including central nervous development (**Figure 4**). Of particular interest, two of the genes included in the central nervous development GO term, *Nr4a2* and *Lmx1b*, are known to be critical for dopaminergic neuron maintenance and development (Luo, 2012; Doucet-Beaupré *et al*., 2016). *Nr4a2* encodes the nuclear receptor related-1 (Nurr1) protein. Nurr1 is a transcription factor involved in dopaminergic neuron development, and it has been suggested that dysregulation of the *Nr4a2* gene may contribute to the pathogenesis of PD (Luo, 2012; Dong *et al*., 2016; Decressac *et al*., 2013). Similarly, *Lmx1b* encodes LIM homeobox transcription factor 1 beta, a transcription factor involved in maintenance of cellular respiration in midbrain dopaminergic neurons (Doucet-Beaupré *et al*., 2016). *Lmx1b* inactivation has been shown to produce Parkinson’s-like cellular features, including a-synuclein inclusions and progressive degeneration of dopaminergic neurons (Doucet-Beaupré *et al*., 2016). Furthermore, while little research has explored the role of human *LMX1B* in Parkinson’s disease, some preliminary evidence suggests a positive association between eleven *LMX1B* polymorphisms and PD in human females (Bergman *et al*., 2009). Given the documented roles of *Nr4a2* and *Lmx1b* in the maintenance and development of dopaminergic neurons, altered regulation of the Nurr1 and Lmx1b transcription factors could influence susceptibility to dopaminergic toxicity. However, the female-specific epigenetic results presented here are difficult to reconcile with previous work showing a male-specific phenotype (Richardson *et al*., 2006). As such, further work must investigate the functional effects of these dieldrin-induced epigenetic effects in females.

In addition to *Nr4a2* and *Lmx1b*, previous work has also identified the related *Pitx3* locus as a potential risk factor for Parkinson’s disease (Li *et al*., 2009). *Pitx3* encodes the pituitary homeobox 3 protein, a transcription factor that shows restricted, constitutive expression in the midbrain, where it is thought to play a role in development and maintenance of dopaminergic neurons (Li *et al*., 2009). Although some studies have shown associations between human *PITX3* polymorphisms and PD (Fuchs *et al*., 2009; Bergman *et al*., 2010; Haubenberger *et al*., 2011), findings have been inconsistent, and a recent meta-analysis suggested that the identified polymorphisms do not show significant associations with risk of PD (Jiménez-Jiménez *et al*., 2014). Reflecting this uncertainty, unlike the *Nr4a2* and *Lmx1b* genes, we did not identify dieldrin-related differential methylation at the *Pitx3* gene (**Figure 5C**). There were also no significant effects of developmental dieldrin exposure on *Pitx3* expression in male or female midbrains (**Figure 6**). These data match previous work showing no change in *Pitx3* expression after developmental exposure to 0.3 mg/kg dieldrin (Richardson *et al*., 2006), suggesting that *Pitx3* regulation is not altered by developmental dieldrin exposure.

### 4.3 Dieldrin-related differential methylation at the imprinted *Grbl0* locus

Imprinted genes, which display parent-of-origin-specific monoallelic expression, are regulated by differentially methylated regions and display particular sensitivity to developmental exposures (Plasschaert and Bartolomei, 2014; Marsit, 2015). Despite these characteristics, we did not identify widespread differential methylation at imprinted genes; however, we did find dieldrin-related hypomethylation in male mice at the imprinted *Grb10* locus (**Figure 5B**). Supporting the functional relevance of these epigenetic findings, the identified *Grb10* DMR was annotated to a predicted mouse midbrain enhancer region, and two protein-coding *Grb10* transcripts demonstrated dieldrin-related changes in expression in the male midbrains (**Figure 6**). *Grb10* encodes growth-factor receptor bound protein 10 (Grb10), a signal adapter protein that regulates cellular proliferation, insulin signaling, and normal adult behavior (Plasschaert and Bartolomei, 2015; Dufresne and Smith, 2005). *Grb10* displays neuron-specific imprinting controlled by differential methylation at an imprinting control region (Plasschaert and Bartolomei, 2015), and the Grb10 protein directly interacts with Grb10-interacting GYF Protein 2 (GIGYF2), a protein encoded within the PARK11 locus (Giovannone *et al*., 2009). Previous work has shown that heterozygous *Gigyf2*^+/-^ mice develop adult-onset neurodegeneration (Giovannone *et al*., 2009), and meta-analyses of GIGYF2 genetic studies have suggested the protein may a role in PD-related neurodegeneration (Lautier *et al*., 2008; Zhang *et al*., 2015). As such, dieldrin-related changes in DNA methylation at *Grb10* have the potential to indirectly impact PD risk by regulating activity of GIGYF2. Future work should investigate whether dieldrin-induced differential methylation at *Grb10* has functional consequences for expression and/or function of Grb10 and GIGYF2.

### 4.4 Poised epigenetic state

Dieldrin-related epigenome-wide differential DNA methylation results were not reflected in genome-wide RNA-sequencing analysis, which completely lacked significance when corrected for multiple testing. Although seemingly contradictory, these data are consistent with the concept of “silent neurotoxicity” and suggest that developmental dieldrin exposure induces a poised methylome that is primed to respond to additional later-life exposures, but does not directly produce changes in phenotype. Such a model fits into the developmental origins of health and disease (DOHaD) paradigm, which holds that early-life environmental exposures modify disease risk into adulthood (Heindel and Vandenberg, 2015). Under this paradigm, the epigenome is a mechanism by which gene regulation is developmentally programmed to respond to later-life perturbations (Bianco-Miotto *et al*., 2017; Gluckman *et al*., 2008). When considering our results from a DOHaD perspective, it makes sense that a developmental exposure could produce clear epigenetic changes without associated changes in gene expression. This work suggests that there is value in novel experimental paradigms that test whether developmental exposures alter sensitivity to toxic insults in adulthood.

### 4.5 Differences between adult and developmental exposures

In this study, there was no overlap between the DMCs and DMRs detected in the adult and developmental exposure models, and the largest effects of dieldrin exposure were seen in the female developmental exposure. Supporting this data, previous studies have shown that the developing brain is particularly sensitive to environmental perturbations (Rauh and Margolis, 2016; Rice and Barone, 2000), and that environmental chemicals can act as potent neurotoxicants during development (Miodovnik, 2011; Heyer and Meredith, 2017). Furthermore, developmental exposures have the potential to modify epigenetic programming, which takes place during embryonic development (Reik *et al*., 2001; Feng *et al*., 2010; Marsit, 2015). These considerations, when combined with our data, suggest that timing of exposure is critical in determining the effects of dieldrin on the midbrain methylome, and that the developing brain is particularly vulnerable to environmental exposures.

### 4.6 Limitations

Although pathway and network analyses provided some context for our identified DMCs, the biological significance of dieldrin-related changes in epigenome-wide DNA methylation remain largely unclear, and future studies will explore the functional relevance of these findings. Since existing gene ontology databases only provide information on documented gene functions, it is quite possible that interpretation of our data is incomplete. Additionally, the RRBS method is inherently biased towards CpG-dense regions, and only covered approximately 3,000,000 CpGs per sample. As a result, we may be missing out on additional dieldrin-related differential methylation in CpG-poor regions. This bias in the RRBS method is particularly concerning for the field of neuroepigenetics, where there is a growing recognition of the importance of DNA methylation along gene bodies, as well as in non-CpG dinucelotides (Lister *et al*., 2013; Guo *et al*., 2014). These issues could be addressed through the use of whole-genome bisulfite sequencing (WGBS-Seq), the gold standard for measuring genome-wide DNA methylation; however, WGBS remains cost prohibitive. Despite these limitations, to our knowledge, this is the first study to show effects of developmental dieldrin exposure on the mouse midbrain methylome.

## Supporting information

## Acknowledgements

The authors thank Marie Adams at the Van Andel Genomics Core for providing consultation, library preparation (RNA-seq), and next-generation sequencing facilities and services (RRBS and RNA-seq). We also thank Claudia Lalancette at the University of Michigan Epigenomics Core for library preparation (RRBS) services.

