## Supplementary material for "Developmental Dieldrin Exposure Alters DNA Methylation at Genes Related to Dopaminergic Neuron Development and Parkinson’s Disease in Mouse Midbrain": RNAseq_DTranscriptE_code_Kochmanski_121018.R

```
######################### START IN LINUX TERMINAL ######################################################

## First step is to quantify RNA-seq data using salmon (https://combine-lab.github.io/salmon/getting_started/)

#Note: Salmon runs in the linux terminal.

#Install salmon using bioconda:

#Download miniconda from https://conda.io/miniconda.html. To install, run the following code (in the directory where the .sh file is located):
bash Miniconda2-latest-MacOSX-x86_64.sh

#Now that miniconda is installed, build channels (including bioconda) for conda database:
conda config --add channels conda-forge
conda config --add channels bioconda
#Also create a channel for salmon:
conda create -n salmon salmon
# To activate this built environment, use:
source activate salmon

#Test to see if salmon is running correctly:
salmon -h   

#Next, we’re going to build an index on our transcriptome. The index is a #structure that salmon uses to quasi-map RNA-seq reads during #quantification. The index need only be constructed once per #transcriptome, and it can then be reused to quantify many experiments. #We use the index command of salmon to build our index:

salmon index -t /path/to/Mus_musculus.GRCm38.cdna.all.fa.gz -i /path/to/mm10_index

#Sample data is collected in three separate flowcell directories. Need to combine the raw flowcell data for each sample prior to use in Salmon.

#Combining multiple flowcell fq.gz files for the same RNA-seq samples -- concatenate files:
cat file1.gz file2.gz file3.gz > allfiles.gz #Example code!

#Run code on actual RNA-seq files
cat /path/to/flowcell1/1_L000_R1_001.fastq.gz /path/to/1_L000_R1_001.fastq.gz /path/to/flowcell3/1_L000_R1_001.fastq.gz  > /path/to/1_L000_R1_combined.fastq.gz

#Repeat the above code for each sample ID - ID #1,2,3,4,5,6,7,8,9,10,11,12,22,23,24,25,26,27,28,29,30,31,32,33

#Note: Also need to run the code on the R2 paired sequencing files -- e.g. 1_L000_R2_001.fastq.gz

#Now that all the sequencing data is in a concatenated gorm, we can quantify our read counts for the sample data.
#Example code from salmon documentation:

salmon quant -i athal_index -l A \
-1 ${fn}/${samp}_1.fastq.gz \
-2 ${fn}/${samp}_2.fastq.gz \
-p 4 -o quants/${samp}_quant

#NOTES: The -i argument tells salmon where to find the index -l A tells salmon #that it should automatically determine the library type of the sequencing #reads (e.g. stranded vs. unstranded etc.). The -1 and -2 arguments tell #salmon where to find the left and right reads for this sample (notice, #salmon will accept gzipped FASTQ files directly). Finally, the -p 8 #argument tells salmon to make use of 8 threads and the -o argument #specifies the directory where salmon’s quantification results sould be #written.

#Example code:
salmon quant -i mm10_index -l A -1 1_L000_R1_001.fastq -2 1_L000_R2_001.fastq.gz -p 4 -o quants/samp1_quant

#Note: Salmon runs in linux, and is too computationally intensive to run on a local computer, so the following steps were run on a cluster.

#Running salmon on the cluster:
ssh -X #Example cluster

#load the installed Salmon module
module load Salmon/0.11.3 

#Linux terminal code for real data:
salmon quant -i /path/to/mm10_index -l A -1 /path/to/1_L000_R1_combined.fastq.gz -2 /path/to/1_L000_R2_combined.fastq.gz -p 8 -o /path/to/samp1_quant
#Repeat for all 24 samples!

########################### END OF LINUX #################################################

#For preparing salmon output for use with sleuth, use the wasabi R package (https://github.com/COMBINE-lab/wasabi).

#The following steps are run in local RStudio console.

#With devtools, it's easy to install wasabi in R:
source("http://bioconductor.org/biocLite.R")
biocLite("devtools")    # only if devtools not yet installed
biocLite("COMBINE-lab/wasabi")

#Once wasabi is installed, you can load the library with:

library(wasabi)

#Now that wasabi is installed, we can use it to convert the Sailfish / #Salmon output into sleuth-compatible format. 
#Imagine we have some samples #(Sailfish quant directories) sitting in a data directory:
#data/samp1_quant   data/samp2_quant   data/samp3_quant   data/samp4_quant

#For our data, the path is:
/path/to/quants/data/samp1_quant
/path/to/quants/data/samp2_quant
/path/to/quants/data/samp3_quant
/path/to/quants/data/samp4_quant
#Etc.

#First, we create a simple vector containing these directories:

sfdirs <- file.path("/path/to/quants/", c("samp1_quant", "samp2_quant", "samp3_quant", "samp4_quant", "samp5_quant", "samp6_quant","samp7_quant", "samp8_quant", "samp9_quant","samp10_quant", "samp11_quant", "samp12_quant","samp22_quant", "samp23_quant", "samp24_quant","samp25_quant", "samp26_quant", "samp27_quant","samp28_quant", "samp29_quant", "samp30_quant","samp31_quant", "samp32_quant", "samp33_quant"))

#Now, we simply run the prepare_fish_for_sleuth function:

prepare_fish_for_sleuth(sfdirs)

#The function will write some status messages to the console and, when it's #done, each directory will now contain and abundance.h5 file in a sleuth-#compatible format. 
#From this point forward, you can simply run sleuth #normally.

#Load libraries:
source("https://bioconductor.org/biocLite.R")
biocLite("tximport")
biocLite("devtools")  
biocLite("remotes")# only if devtools not yet installed
biocLite("COMBINE-lab/wasabi")
biocLite("pachterlab/sleuth")
install.packages("cowplot")
biocLite("goseq") #Includes dependency - 'geneLenDataBase'
library(tximport)
library(readr)
library(tidyr)
library(plyr)
library(devtools)
library(wasabi)
library(sleuth)
library(cowplot)
library(goseq)
library(biomaRt)

#==============================
# divide into male & female datasets
sex.id.M = 'M' # male data
sex.id.F = 'F' #female data
meta.M = meta[which(meta$sex==sex.id.M),]
meta.F = meta[which(meta$sex==sex.id.F),]
#==============================

#Remove files in the working dir
#Get the sample ids
sample_id <- dir(file.path("/path/to/directory/", "data_sleuth"))
sample_id #lists sample ids.

# Note: File directory path variable needs to be in the sample_to_covariates (meta) df for sleuth_prep
#Read in table of meta data that includes /path for each sample's wasabi-processed data.
meta.path <- read.table("/path/to/directory/P3_RNAseq_Meta_Wasabi_Path_100318.txt", header = TRUE)
print(meta.path)

#stratify by sex
meta.path.F <- read.table("/path/to/directory/P3_RNAseq_Meta_Wasabi_Path_Female_100318.txt", header = TRUE)
print(meta.path.F)

meta.path.M <- read.table("/path/to/directory/P3_RNAseq_Meta_Wasabi_Path_Male_100318.txt", header = TRUE)
print(meta.path.M)

#Side note: path must be character for running in sleuth:
meta.path$path <- as.character(meta.path$path)
meta.path$path

meta.path.F$path <- as.character(meta.path.F$path)
meta.path.F$path
meta.path.M$path <- as.character(meta.path.M$path)
meta.path.M$path


#############################################################################################

#Build databases

# Set the host
# mart <- biomaRt::useMart(biomart = 'ENSEMBL_MART_ENSEMBL')
# mart <- biomaRt::useMart(biomart = 'ENSEMBL_MART_ENSEMBL'); View(listDatasets(mart))
mart <- biomaRt::useMart(biomart = 'ENSEMBL_MART_ENSEMBL', host="useast.ensembl.org",dataset = 'mmusculus_gene_ensembl') # this is the dataset (Mouse genes (GRCm38.p5) used by UCSC to create their mm10 database

t2g <- biomaRt::getBM(attributes = c("ensembl_transcript_id", "ensembl_gene_id", "external_gene_name", "description"), mart = mart)

t2g <- dplyr::rename(t2g, ens_transcr = ensembl_transcript_id, ens_gene = ensembl_gene_id, ext_gene = external_gene_name, predicted_function = description)

# Rename again for Sleuth
t2gSleuth <- dplyr::rename(t2g, target_id = ens_transcr) # target is the transcript

############# DTE with Sleuth ##########################################

#Sleuth prep
full_design <- model.matrix(~meta.path$treatment) # NO COVARIATES; ONLY TREATMENT.
full_design.F <- model.matrix(~meta.path.F$treatment) # NO COVARIATES; ONLY TREATMENT.
full_design.M <- model.matrix(~meta.path.M$treatment) # NO COVARIATES; ONLY TREATMENT.

colnames(full_design)[1] <- c("Intercept") #Create variable for intercept.
colnames(full_design.F)[1] <- c("Intercept") #Create variable for intercept.
colnames(full_design.M)[1] <- c("Intercept") #Create variable for intercept.

#A basic filter to apply to sleuth prep -- 50% of samples must have at least 10 est counts
my_basic_filter <- function(row, min_reads = 10, min_prop = 0.5) {
  mean(row >= min_reads) >= min_prop
}

#The 'sleuth' object stores a group of wasabi results, and can then operate on them while accounting for covariates, sequencing depth, technical and biological variance.
so <- sleuth_prep(meta.path, full_model = full_design, target_mapping = t2gSleuth, filter_fun = my_basic_filter, extra_bootstrap_summary = TRUE)

#Stratify models by sex:
so.F <- sleuth_prep(meta.path.F, full_model = full_design.F, target_mapping = t2gSleuth, filter_fun = my_basic_filter, extra_bootstrap_summary = TRUE)
so.M <- sleuth_prep(meta.path.M, full_model = full_design.M, target_mapping = t2gSleuth, filter_fun = my_basic_filter, extra_bootstrap_summary = TRUE)

#Fit the models
so <- sleuth_fit(so, fit_name="full") # full model, uses full_design
so.F <- sleuth_fit(so.F, fit_name="full") # full model, uses full_design to design model based on treatment.
so.M <- sleuth_fit(so.M, fit_name="full") # full model, uses full_design to design model based on treatment.

#What this has accomplished is to “smooth” the raw kallisto abundance estimates for each sample using a linear model with a parameter that represents the experimental condition. 
#To test for transcripts that are differential expressed between the conditions, sleuth performs a second fit to a “reduced” model that presumes abundances are equal in the two conditions. 
#To identify differential expressed transcripts sleuth will then identify transcripts with a significantly better fit in the “full” model.
#Fit the reduced model(s). In this case the 'reduced' model is one with ONLY the intercept (null model):
so <- sleuth_fit(so, formula = ~1, fit_name = "reduced")
so.F <- sleuth_fit(so.F, formula = ~1, fit_name = "reduced")
so.M <- sleuth_fit(so.M, formula = ~1, fit_name = "reduced")

#View the models
models(so)
models(so.F)
models(so.M)

#Likelihood ratio test (DIFFERENTIAL TRANSCRIPT EXPRESSION):
so <- sleuth_lrt(so, "reduced", "full")
so.F <- sleuth_lrt(so.F, "reduced", "full")
so.M <- sleuth_lrt(so.M, "reduced", "full")

#Gene-level differential expression
#When running the command ‘sleuth_results,’ sleuth uses the p-values from comparing transcripts to make a gene-level determination and perform gene differential expression.
sleuth_table_gene.F <- sleuth_results(so.F, 'reduced:full', 'lrt', show_all = FALSE) 
sleuth_table_gene.F.sig <- dplyr::filter(sleuth_table_gene.F, qval <= 0.10) # 0 observations
sleuth_table_gene.M <- sleuth_results(so.M, 'reduced:full', 'lrt', show_all = FALSE) 
sleuth_table_gene.M.sig <- dplyr::filter(sleuth_table_gene.M, qval <= 0.10) # 0 observations 

#The most significant genes are pulled out using the following command:
head(sleuth_table_gene.F, 20)
head(sleuth_table_gene.M, 20)

sleuth_table <- sleuth_results(so, test='reduced:full', test_type='lrt', show_all = FALSE)
sleuth_significant <- dplyr::filter(sleuth_table, qval <= 0.05) #0 observations w/ q-value < 0.05
sleuth_significant <- dplyr::filter(sleuth_table, qval <= 0.10) #0 observations

#DTE lists
#Female only:
sleuth_table.F <- sleuth_results(so.F, test='reduced:full', test_type='lrt', show_all = FALSE)
sleuth_significant.F <- dplyr::filter(sleuth_table.F, qval <= 0.05) #0 observations w/ q-value < 0.05
sleuth_significant.F <- dplyr::filter(sleuth_table.F, qval <= 0.10) #0 observations
#Male only:
sleuth_table.M <- sleuth_results(so.M, test='reduced:full', test_type='lrt', show_all = FALSE)
sleuth_significant.M <- dplyr::filter(sleuth_table.M, qval <= 0.05) #0 observations w/ q-value < 0.05 
sleuth_significant.M <- dplyr::filter(sleuth_table.M, qval <= 0.10) #0 observations

#Write the significant results to a .csv (Note: didn't actually do this due to lack of significance):
write.csv(sleuth_significant, "/path/to/directory/RNAseq_Sleuth_DTE_Sig.csv") # 0 observations
#Write sleuth test tables (including non-significant results) to .csv files:
write.csv(sleuth_table.M, "/path/to/directory/RNAseq_Sleuth_DTE_Male_TestTable.csv")
write.csv(sleuth_table.F, "/path/to/directory/RNAseq_Sleuth_DTE_Female_TestTable.csv")

#go to sleuth live to view results in tabular form; useful R shiny interface for examining specific genes/transcripts.
sleuth_live(so)
sleuth_live(so.F)
sleuth_live(so.M)

#Turn off sleuth_live interface.
dev.off()

#############################################################################################

## Create full DTE plots (w/ both sexes) for gene transcripts.

install.packages("ggsignif")
library(ggsignif)

#Plot the Nr4a2/Nurr1 transcript with marginally significant differential expression in the female mice (transcript 201):
#Plot transcript Nr4a2-201 (p=0.16 in males; p=0.06 in females).
#Plot both male and female data:
Nurr1_DTE = read.csv("/path/to/directory/P3_RNAseq_Nurr1_Transcript201_Raw-Est-Counts_102218.csv")

#Reorder sex variable for plotting purposes:
Nurr1_DTE$sex_1 = factor(Nurr1_DTE$sex, levels=c('Male','Female'))

ggplot(Nurr1_DTE, aes(x = treatment, y = est_counts, color = treatment)) +
  geom_jitter(position = position_jitter(width = 0.05, height = 0), size = 3) +
  stat_summary(fun.y = median, fun.ymin = median, fun.ymax = median,
               geom = "crossbar", width = 0.5) +
  ylim(200, 1200) +
  scale_color_manual(values=c("#008183", "#F08521")) + #change colors
  facet_grid(cols = vars(sex_1)) + #grid by sex
  labs(x = "Treatment", y = "Estimated counts (n)") + #change axis labels 
  annotate("text", x = "Dieldrin", y = 1100, label = c(" ", "*"), size = 8.5) #add asterisk for significance
```
