## Supplementary material for "Developmental Dieldrin Exposure Alters DNA Methylation at Genes Related to Dopaminergic Neuron Development and Parkinson’s Disease in Mouse Midbrain": RRBS_adult exposure_differential_methyl_code_Kochmanski_121018.Rmd

````
---
title: "P2_RRBS_complete_differential_meth_112818"
output: html_document
---

```{r setup, include=FALSE}
knitr::opts_chunk$set(echo = TRUE)
```

## R Markdown

---
title: "DSS_RRBS Differential Methylation_112818"
author: "Joe Kochmanski"
date: "November 28, 2018"
output:
  html_document:
    code_folding: hide
    fig_height: 2.5
    fig_width: 3.5
    highlight: tango
    toc: yes
    toc_float: yes
---

## Project summary

This script performs differential methylation analysis from RRBS data.

1. Read in bismark coverage files using bsseq. 
2. DSS Package - differential methylation modeling for RRBS data.
3. DMR calling on adult exposure-related DMCs.
4. Write CSVs for results from models sorted by p-values.
5. Examine the test statistics and p-values for the female and male datasets.
6. Annotate DMCs/DMRs using annotatr.

## Load required packages        
```{r setup, echo=TRUE, warning=FALSE, message=FALSE, cache=FALSE, results='hide'}
source("https://bioconductor.org/biocLite.R")
biocLite("bsseq")
biocLite("DSS")
biocLite("annotatr")
biocLite("DMRcate")
library(bsseq)
library(DSS)
library(annotatr)
library(DMRcate)
```

## 1. Read in bismark coverage files using bsseq
```{r Read coverage data, echo=TRUE, warning=FALSE, message=FALSE, cache=FALSE}

##Read in .cov bismark files for each sample
##Example:
#read.bismark(files, sampleNames, rmZeroCov = TRUE, strandCollapse = TRUE, fileType = c("cov", "oldBedGraph", "cytosineReport"), mc.cores = 1, verbose = TRUE)

################ Read in the RRBS coverage data ##################

setwd("/path/to/directory/")

###Read in RRBS data for adult male exposure study (only one sex, so no stratification):
bsseq_P2 <- read.bismark(files = c("P2-E1-005MockBS_L000_R1_001_val_1_bismark_bt2_pe.bismark.cov.gz",#C
                                        "P2-E1-007MockBS_L000_R1_001_val_1_bismark_bt2_pe.bismark.cov.gz",#C
                                        "P2-E1-009MockBS_L000_R1_001_val_1_bismark_bt2_pe.bismark.cov.gz",#C
                                        "P2-E1-010MockBS_L000_R1_001_val_1_bismark_bt2_pe.bismark.cov.gz",#C
                                        "P2-E1-013MockBS_L000_R1_001_val_1_bismark_bt2_pe.bismark.cov.gz",#C
                                        "P2-E1-002MockBS_L000_R1_001_val_1_bismark_bt2_pe.bismark.cov.gz",#D
                                        "P2-E1-014MockBS_L000_R1_001_val_1_bismark_bt2_pe.bismark.cov.gz",#D
                                        "P2-E1-015MockBS_L000_R1_001_val_1_bismark_bt2_pe.bismark.cov.gz",#D
                                        "P2-E1-017MockBS_L000_R1_001_val_1_bismark_bt2_pe.bismark.cov.gz",#D
                                        "P2-E1-028MockBS_L000_R1_001_val_1_bismark_bt2_pe.bismark.cov.gz"),#D
sampleNames = c("C_1","C_2","C_3","C_4","C_5","D_1","D_2","D_3","D_4", "D_5"),
                               rmZeroCov = TRUE,
                               strandCollapse = FALSE,
                               fileType = "cov",
                               verbose = TRUE)

```


## 2. DSS Package - differential methylation modeling for RRBS data
```{r Establish model design, echo=TRUE, warning=FALSE, message=FALSE, cache=FALSE}

#Stratify models by sex when examining exposure effects in mouse RRBS data:
DMLtest_P2.exp = DMLtest(bsseq_P2, group1=c("C_1","C_2","C_3","C_4","C_5"), 
                               group2=c("D_1","D_2","D_3","D_4", "D_5"), 
                               equal.disp = FALSE, smoothing = FALSE)

#Smoothing = FALSE for RRBS data (non-continuous coverage)

#View top few results for each analysis:
head(DMLtest_P2.exp)

#Call DMLs from test results:
DMLtest_P2.exp

##Sort the dieldrin-related DMCs

#Sort the data frames by p-value (and remove NAs) using following codes:
DMLtest_P2.exp_sort1 <- DMLtest_P2.exp[order(DMLtest_P2.exp$pval, na.last=NA), ]

head(DMLtest_P2.exp_sort1) #View first few significant results

#Subset data frame based on FDR cutoff of 0.05
DMLtest_P2.exp_sort2 <- subset(DMLtest_P2.exp_sort1, fdr < 0.05)
head(DMLtest_P2.exp_sort2) #View first few significant results
summary(DMLtest_P2.exp_sort2) #View summary of data frame variables

#Institute more stringent cutoffs for DMC-calling -- at least 5% change in methylation by exposure cutoff
DMLtest_P2.exp_sort3 <- subset(DMLtest_P2.exp_sort2, abs(diff) > 0.05)
#Note: None of the DMCs with FDRs<0.05 were removed, indicating that all of these had an absolute difference of >5% methylation.

#99 DMCs by dieldrin exposure.

head(DMLtest_P2.exp_sort3) #View first few significant results
```


## 3.DMR calling on adult exposure-related DMCs
```{r DMR calling on called DMCs, echo=TRUE, warning=FALSE, message=FALSE, cache=FALSE}
dmrs_P2.exp1 <- callDMR(DMLtest_P2.exp_sort1, p.threshold=0.01, minlen=20, minCG=3) #  3 DMRs
dmrs_P2.exp2 <- callDMR(DMLtest_P2.exp_sort2, p.threshold=0.01, minlen=20, minCG=3) # 0 DMRs
dmrs_P2.exp3 <- callDMR(DMLtest_P2.exp_sort3, p.threshold=0.01, minlen=20, minCG=3) # 0 DMRs
#Can manually adjust the p-value threshold for DMR calling.
dmrs_P2.exp4 <- callDMR(DMLtest_P2.exp_sort1, p.threshold=0.05, minlen=20, minCG=3) # 8 DMRs
dmrs_P2.exp5 <- callDMR(DMLtest_P2.exp_sort2, p.threshold=0.05, minlen=20, minCG=3) # 0 DMRs
dmrs_P2.exp6 <- callDMR(DMLtest_P2.exp_sort3, p.threshold=0.05, minlen=20, minCG=3) # 0 DMRs
#Can also manually adjust the length threshold for DMR calling.
dmrs_P2.exp7 <- callDMR(DMLtest_P2.exp_sort1, p.threshold=0.05, minlen=10, minCG=3) # 9 DMRs
dmrs_P2.exp8 <- callDMR(DMLtest_P2.exp_sort2, p.threshold=0.05, minlen=10, minCG=3) # 0 DMRs
dmrs_P2.exp9 <- callDMR(DMLtest_P2.exp_sort3, p.threshold=0.05, minlen=10, minCG=3) # 0 DMRs

#Notes: Different p-value thresholds were attempted. For final DMR calling, minimum length was set to 10 bp; minimum # of CpGs was set to 3.

#################################################################################################
#DMR calling from DSS gave only a few DMRs, so try more flexible DMR calling function -- DMRcate:
source("https://bioconductor.org/biocLite.R")
biocLite("DMRcate")
library(DMRcate)

wgbsannot.P2 <- cpg.annotate("sequencing", DMLtest_P2.exp)

detectCores() #Use this to determine whether parallel processing is okay to attempt. If not, stick with mc.cores =1.

#Notes about DMRcate: The values of lambda and C should be chosen with care. For array data, the DMRcate authors currently #recommend that half a kilobase represents 1 standard deviation of support (lambda=1000 and C=2), and 20bp
#(C=50) for WGBS data. For RRBS data, we tried a variety of parameters, settling on the settings below:

#Multiple settings used to call DMRs using DMRcate:
#Try restoring p-value cutoff to default (as recommened in DMRcate documentation):
wgbs.P2.DMRs2 <- dmrcate(wgbsannot.P2, lambda = 200, C = 10, min.cpgs = 3, mc.cores = 1)
wgbs.P2.DMR.ranges2 <- extractRanges(wgbs.P2.DMRs2, genome = "mm10")
P2_DMRs_DMRcate2 <- as.data.frame(wgbs.P2.DMR.ranges2) #1 DMR w/ Stouffer < 0.1

#Try adjusting lambda and C values:
wgbs.P2.DMRs3 <- dmrcate(wgbsannot.P2, lambda = 100, C = 20, min.cpgs = 3, mc.cores = 1)
wgbs.P2.DMR.ranges3 <- extractRanges(wgbs.P2.DMRs3, genome = "mm10")
P2_DMRs_DMRcate3 <- as.data.frame(wgbs.P2.DMR.ranges3) #0 DMRs w/ Stouffer < 0.1

#Go higher with the lambda value
wgbs.P2.DMRs4 <- dmrcate(wgbsannot.P2, lambda = 500, C = 4, min.cpgs = 3, mc.cores = 1)
wgbs.P2.DMR.ranges4 <- extractRanges(wgbs.P2.DMRs4, genome = "mm10")
P2_DMRs_DMRcate4 <- as.data.frame(wgbs.P2.DMR.ranges4) #0 DMRs w/ Stouffer < 0.1

#Only the first attempt produced any significance, so those settings were used for the final DMR calling.
setwd("/path/to/directory/")
write.csv(P2_DMRs_DMRcate2, file = "DSS_DMRs_P2_RRBS_Dieldrin_DMRcate_091818.csv")

```

## 4. Write CSVs for results from models sorted by p-values
```{r DMR calling on called DMCs, echo=TRUE, warning=FALSE, message=FALSE, cache=FALSE}
setwd("/path/to/directory/")
write.csv(DMLtest_P2.exp_sort3, file = "DSS_DMCs_P2_RRBS_Dieldrin_091718.csv")
write.csv(dmrs_P2.exp7, file = "DSS_DMRs_P2_RRBS_Dieldrin_091718.csv")
```

## 5.Examine the test statistics and p-values for the female and male datasets
```{r P values and test statistics, echo=TRUE, warning=FALSE, message=FALSE, cache=FALSE}
par(mfrow=c(1,2))
hist(DMLtest_P2.exp_sort3$stat, 50, main="Combined Exposure test statistics", xlab="")
hist(DMLtest_P2.exp_sort3$pval, 50, main="Combined Exposure P-values", xlab="")
hist(DMLtest_P2.exp_sort3$fdr, 50, main="Combined Exposure FDRs", xlab="")

```

## 6. Annotate DMCs/DMRs using annotatr
```{r P values and test statistics, echo=TRUE, warning=FALSE, message=FALSE, cache=FALSE}
###############BED format:
#In excel, remove top row of variable names and fix chromosome numbers, then save file as .txt file. File should be in standard BED3 format plus any additional columns of sample information.
#This file can then be read in by annotatr to annotate gene regions

##Annotate CpG sites using the annotatr package
source("https://bioconductor.org/biocLite.R")
biocLite("annotatr")
install.packages("stringi")
biocLite("regioneR")
biocLite("TxDb.Mmusculus.UCSC.mm10.knownGene")
biocLite("org.Mm.eg.db")
library(annotatr)
library(rtracklayer)
library(regioneR)
library(TxDb.Mmusculus.UCSC.mm10.knownGene)
library(org.Mm.eg.db)

setwd("C:/")
setwd("/path/to/directory")

################# Adding CUSTOM miRNA and enhancer annotations to annotatr #################

library(GenomicRanges)
library(rtracklayer)
library(annotatr)

setwd("/path/to/directory/")
mirna_gff <- import.gff("mmu_miRNA_annotations.gff3") # Imports mus musculus miRNA GFF3 genome annotation file (from: http://www.mirbase.org/ftp.shtml).
mirna_gff[1:4,]

#Coerce miRNA database to a data frame in R
mirna_gff_df <- as.data.frame(mirna_gff)

#Write .csv of data frame for conversion into BED6 format in excel:
setwd("/path/to/directory/")
write.csv(mirna_gff_df, "miRNA_database_annotations_081418.csv")

## Custom annotation objects are given names of the form genome_custom_name
setwd("/path/to/directory/")
mirna_db_file = "miRNA_database_annotations_081418_BEDformat.txt"
extraCols0 = c(gene_id='character', symbol='character', type='character')
#Use the read_annotations function to create custom annotation file for miRNA:
read_annotations(con = mirna_db_file, genome = 'mm10', name = 'miRNA',  extraCols = extraCols0, format='bed')

#Also create custom annotation for predicted enhancers identified from postnatal mouse midbrain (ENCODE).
#Raw file downloaded from: https://www.encodeproject.org/annotations/ENCSR114ZIJ/
setwd("/path/to/directory/")
enhancer_db_file = "ENCFF129YCY_ENCODE_mouse_midbrain_enhancers_091318.bed"
extraCols0 = c(gene_id='character', symbol='character', type='character')
read_annotations(con = enhancer_db_file, genome = 'mm10', name = 'enhancer',  extraCols = extraCols0, format='bed')

#Double-check that custom annotation track has been successfully added to the mm10
print(annotatr_cache$get('mm10_custom_miRNA')) #8/14/18 -- it works!
print(annotatr_cache$get('mm10_custom_enhancer')) #9/13/18 -- it works!

##################### Annotating REAL DATA ##############################

#Note for annotatr: Used BED6+3 format file in the following format:
#chr4	  152512417	152512418	hyper	-0.953057058	0.033368133	0.986425191
#chr12	48140364	48140365	hyper	-0.932408194	0.044534154	0.976942348

#Columns four, five, six, and seven refer to direction of change, magnitude of change (diff),
#mu1, and mu2, respectively.

#Note that the start column is set to -1 compared to the end column; this 
#ensures that the final form of the data will have identical start and end values
#aftering reading in the data using annotatr.

#Set working directory and read in bed file
setwd("/path/to/directory/")
dmc_P2_file = "DSS_DMCs_P2_RRBS_Dieldrin_BEDformat_091718.txt"

file.exists("/Users/JoeKochmanski/Dropbox/Bernstein_Lab/Sequencing_Data_Analysis/DSS_DM_Output/P2_DSS_DMCs/DSS_DMCs_P2_RRBS_Dieldrin_BEDformat_091718.txt")


#Define and rename extra columns in BED file:
extraCols = c(diff_meth = 'numeric', mu1 = 'numeric', mu2 = 'numeric')
dmc_P2_regions = read_regions(con = dmc_P2_file, extraCols = extraCols, genome = 'mm10',
                              rename_name = 'DM_status', format = 'bed')
head(dmc_P2_regions)

#Build annotations for mm10 based on variable of interest (dieldrin exposure):
builtin_annotations()
annots_exp = c('mm10_basicgenes','mm10_cpgs', 'mm10_lncrna_gencode', 'mm10_custom_miRNA', 'mm10_custom_enhancer')
annotations_exp = build_annotations(genome = 'mm10', annotations = annots_exp)

#Annotate the regions:
dmc_annots_P2_exp = annotate_regions(
  regions = dmc_P2_regions,
  annotations = annotations_exp,
  ignore.strand = TRUE)

# Note: annotatr output is a special type of dataframe using the dplyr package.
# By default, dplyr::tbl_df objects have nice printing properties, but it
# hides extra columns that would ordinarily wrap. You can see them all with:
head(as.data.frame(dmc_annots_P2_exp))

#Write the gene annotations to a tab-delimited text file:
setwd("/path/to/directory/")
write.table(as.data.frame(dmc_annots_P2_exp),file="DSS_DMCs_P2_RRBS_Dieldrin_Annotated_091818.txt",sep="\t",quote=F)

##Distributions of annotated DMCs

#Define order of differentially methylated CpGs (DMCs)
dm_order = c(
  'hyper',
  'hypo')
cpg_order = c(
  'mm10_cpg_inter',
  'mm10_cpg_islands',
  'mm10_cpg_shores',
  'mm10_cpg_shelves',
  'mm10_genes_promoters',
  'mm10_genes_3UTRs',
  'mm10_genes_5UTRs',
  'mm10_genes_exons',
  'mm10_genes_introns',
  'mm10_lncrna_gencode',
  'mm10_custom_enhancer') #NOTE: Tried including custom miRNA track, but none of CpGs annotated to any miRNAs, so I removed this track from the graphs (it was showing as NA).
#NOTE: Tried including custom enhancer track from ENCODE. There are indeed some CpG sites that map to enhancers!

#Plot categorical to view CpG distributions vs. random distribution.
dmc_P2_exp = plot_categorical(
  annotated_regions = dmc_annots_P2_exp,
  x = 'DM_status',
  fill = 'annot.type',
  x_order = dm_order,
  fill_order = cpg_order,
  position = 'fill',
  legend_title = 'knownGene Annotations',
  x_label = 'DM status',
  y_label = 'Proportion')
plot(dmc_P2_exp) #No DMCs annotated to 3'UTRs.

#Save Exposure-based DMC distribution R plot to ONES server:
setwd("/path/to/directory/")
pdf("DSS_DMCs_P2_RRBS_Dieldrin_Distribution_091818.pdf") 
plot(dmc_P2_exp)
dev.off()

# Create randomized regions for adult exposure study
dmc_rnd_P2_regions = randomize_regions(regions = dmc_P2_regions)
dmc_rnd_P2_annots = annotate_regions(
  regions = dmc_rnd_P2_regions,
  annotations = annotations_exp,
  ignore.strand = TRUE)
dmc_exp_P2_rnd = plot_categorical(
  annotated_regions = dmc_annots_P2_exp,
  annotated_random = dmc_rnd_P2_annots,
  x = 'DM_status',
  fill = 'annot.type',
  x_order = dm_order,
  fill_order = cpg_order,
  position = 'fill',
  legend_title = 'knownGene Annotations',
  x_label = 'DM status',
  y_label = 'Proportion') #No 3'UTRs.

#Plot actual CpGs compared to random distribution.
setwd("/path/to/directory/")
pdf("DSS_DMCs_P2_RRBS_Dieldrin_Distribution+Random_091818.pdf") 
plot(dmc_exp_P2_rnd)
dev.off()

#################### Repeat this process for the DMRs ####################

#Read in DMRs
setwd("/path/to/directory/")

#DSS callDMR() DMRs
dmr_P2_file = "DSS_DMRs_P2_RRBS_Dieldrin_BEDformat_091718.txt"

#Define and rename extra columns in BED file:
extraCols = c(diff_meth = 'numeric', mu1 = 'numeric', mu2 = 'numeric')
dmr_P2_regions = read_regions(con = dmr_P2_file, extraCols = extraCols, genome = 'mm10',
                              rename_name = 'DM_status', format = 'bed')
head(dmr_P2_regions)


#DSS + DMRcate DMRs:
setwd("/path/to/directory/")
dmr_P2_file2 = "DSS_DMRs_P2_RRBS_Dieldrin_DMRcate_BEDformat_091818.txt"
#DMRcate DMRs (BED3 format; No extra columns.)
dmr_P2_regions2 = read_regions(con = dmr_P2_file2, genome = 'mm10',
                               format = 'bed')
head(dmr_P2_regions2)


#Build annotations for mm10 based on variable of interest (age):
annots_exp = c('mm10_basicgenes','mm10_cpgs', 'mm10_lncrna_gencode', 'mm10_custom_miRNA', 'mm10_custom_enhancer')
annotations_exp = build_annotations(genome = 'mm10', annotations = annots_exp)

##Now annotate!

#DSS DMRs:
dmrs_P2_exp_annotated = annotate_regions(regions = dmr_P2_regions, annotations = annotations_exp, ignore.strand = TRUE)

#DMRcate DMRs:
dmrs_P2_exp_annotated2 = annotate_regions(regions = dmr_P2_regions2, annotations = annotations_exp, ignore.strand = TRUE)

# Note: annotatr output is a special type of dataframe using the dplyr package.
# By default, dplyr::tbl_df objects have nice printing properties, but it
# hides extra columns that would ordinarily wrap. You can see them all with:
head(as.data.frame(dmrs_P2_exp_annotated))

## You could write the gene region annotations to a tab-delimited text file with:
setwd("/path/to/directory/")
write.table(as.data.frame(dmrs_P2_exp_annotated),file="DSS_DMRs_P2_RRBS_Dieldrin_Annotated_091818.txt",sep="\t",quote=F)

#DMRcate DMRs:
setwd("/path/to/directory/")
write.table(as.data.frame(dmrs_P2_exp_annotated2),file="DSS_DMRs_P2_RRBS_Dieldrin_DMRcate_Annotated_091818.txt",sep="\t",quote=F)

##Too few DMRs to generate distributions by location.
```
````
