## Supplementary material for "Developmental Dieldrin Exposure Alters DNA Methylation at Genes Related to Dopaminergic Neuron Development and Parkinson’s Disease in Mouse Midbrain": RRBS_developmental exposure_differential_methyl_code_Kochmanski_121018.Rmd

````
---
title: "RRBS Differential Methylation R Code - 11/20/18"
output: html_document
---

```{r setup, include=FALSE}
knitr::opts_chunk$set(echo = TRUE)
```

## R Markdown

---
title: "RRBS Differential Methylation R Code - 11/20/18"
author: "Joe Kochmanski"
date: "November 20, 2018"
output:
  html_document:
    code_folding: hide
    fig_height: 2.5
    fig_width: 3.5
    highlight: tango
    toc: yes
    toc_float: yes
---

## Project summary

This script performs differential methylation analysis from RRBS data.

1. Read in bismark coverage files using bsseq. 
2. DSS Package - differential methylation modeling for RRBS data.
3. DMR calling on exposure-related DMCs.
4. Write CSVs for results from models sorted by p-values.
5. Examine the test statistics and p-values for the female and male datasets.
6. Annotate DMCs/DMRs using annotatr.
7. Visualize identified DMRs.

## Install and load required packages        
```{r setup, echo=TRUE, warning=FALSE, message=FALSE, cache=FALSE, results='hide'}
source("https://bioconductor.org/biocLite.R")
biocLite("bsseq")
biocLite("DSS")
biocLite("annotatr")
biocLite("DMRcate")
library(bsseq)
library(DSS)
library(annotatr)
library(DMRcate)
```

## 1. Read in bismark RRBS coverage data using bsseq
```{r Read coverage data, echo=TRUE, warning=FALSE, message=FALSE, cache=FALSE}

##Read in .cov bismark files for each sample
##Example:
#read.bismark(files, sampleNames, rmZeroCov = TRUE, strandCollapse = TRUE, fileType = c("cov", "oldBedGraph", "cytosineReport"), mc.cores = 1, verbose = TRUE)

#Set working directory
getwd()
setwd("/path/to/working/directory/")

##################### Notes about DSS package #################
#The core of DSS is a new dispersion shrinkage method for estimating the dispersion parameter from Gamma-Poisson or Beta-Binomial distributions.

#The BS-seq count data are modeled as Beta-Binomial distribution, where the biological variations are captured by the dispersion parameter. The dispersion parameters are estimated through a shrinakge estimator based on a Bayesian hierarchical model. Then a Wald test is performed at each CpG site.
#Due to the differences in coverages, some CpG sites have no coverage in one or both groups, and the test cannot be performed. Those loci will be ignored in testing and results will be "NA".

#Note: In DSS, BS-seq data from a general experimental design (such as crossed experiment, or experiment with covariates) can also be modeled through a generalized linear model framework. 

################ Read in the coverage data ##################

setwd("/path/to/Female RRBS Coverage Files")

###Make bsseq file for females:
bsseq_females <- read.bismark(files = c("P3-E1-053MockBS_L000_R1_001_val_1_bismark_bt2_pe.bismark.cov.gz",#Control
                                        "P3-E1-063MockBS_L000_R1_001_val_1_bismark_bt2_pe.bismark.cov.gz",#Control
                                        "P3-E1-073MockBS_L000_R1_001_val_1_bismark_bt2_pe.bismark.cov.gz",#Control
                                        "P3-E1-098MockBS_L000_R1_001_val_1_bismark_bt2_pe.bismark.cov.gz",#Control
                                        "P3-E1-104MockBS_L000_R1_001_val_1_bismark_bt2_pe.bismark.cov.gz",#Control
                                        "P3-E1-121MockBS_L000_R1_001_val_1_bismark_bt2_pe.bismark.cov.gz",#Control
                                        "P3-E1-132MockBS_L000_R1_001_val_1_bismark_bt2_pe.bismark.cov.gz",#Dieldrin
                                        "P3-E1-140MockBS_L000_R1_001_val_1_bismark_bt2_pe.bismark.cov.gz",#Dieldrin
                                        "P3-E1-164MockBS_L000_R1_001_val_1_bismark_bt2_pe.bismark.cov.gz",#Dieldrin
                                        "P3-E1-172MockBS_L000_R1_001_val_1_bismark_bt2_pe.bismark.cov.gz",#Dieldrin
                                        "P3-E1-179MockBS_L000_R1_001_val_1_bismark_bt2_pe.bismark.cov.gz",#Dieldrin
                                       "P3-E1-195MockBS_L000_R1_001_val_1_bismark_bt2_pe.bismark.cov.gz"),#Dieldrin
sampleNames = c("C_1","C_2","C_3","C_4","C_5","C_6","D_1","D_2","D_3","D_4", "D_5", "D_6"),
                               rmZeroCov = TRUE,
                               strandCollapse = FALSE,
                               fileType = "cov",
                               verbose = TRUE)


setwd("/path/to/Male RRBS Coverage Files")

#Also make separate bsseq file for males:                                         
bsseq_males <- read.bismark(files = c("P3-E1-051MockBS_L000_R1_001_val_1_bismark_bt2_pe.bismark.cov.gz",  #C
"P3-E1-116MockBS_L000_R1_001_val_1_bismark_bt2_pe.bismark.cov.gz", #C
"P3-E1-100MockBS_L000_R1_001_val_1_bismark_bt2_pe.bismark.cov.gz", #C
"P3-E1-058MockBS_L000_R1_001_val_1_bismark_bt2_pe.bismark.cov.gz", #C
"P3-E1-065MockBS_L000_R1_001_val_1_bismark_bt2_pe.bismark.cov.gz", #C
"P3-E1-094MockBS_L000_R1_001_val_1_bismark_bt2_pe.bismark.cov.gz", #C
"P3-E1-159MockBS_L000_R1_001_val_1_bismark_bt2_pe.bismark.cov.gz", #D
"P3-E1-167MockBS_L000_R1_001_val_1_bismark_bt2_pe.bismark.cov.gz", #D
"P3-E1-175MockBS_L000_R1_001_val_1_bismark_bt2_pe.bismark.cov.gz", #D
"P3-E1-192MockBS_L000_R1_001_val_1_bismark_bt2_pe.bismark.cov.gz", #D
"P3-E1-127MockBS_L000_R1_001_val_1_bismark_bt2_pe.bismark.cov.gz", #D
"P3-E1-135MockBS_L000_R1_001_val_1_bismark_bt2_pe.bismark.cov.gz"), #D                                  
sampleNames = c("C_1","C_2","C_3","C_4","C_5","C_6","D_1","D_2","D_3","D_4","D_5","D_6"),
                               rmZeroCov = TRUE,
                               strandCollapse = FALSE,
                               fileType = "cov",
                               verbose = TRUE)

```


## 2. DSS Package - differential methylation modeling for RRBS data
```{r Establish model design, echo=TRUE, warning=FALSE, message=FALSE, cache=FALSE}

#Sex-stratified differential methylation models to examining exposure effects in mouse RRBS data:
DMLtest_female.exp = DMLtest(bsseq_females, group1=c("C_1","C_2","C_3","C_4","C_5","C_6"), 
                               group2=c("D_1","D_2","D_3","D_4", "D_5", "D_6"), 
                               equal.disp = FALSE, smoothing = FALSE)

DMLtest_male.exp = DMLtest(bsseq_males, group1=c("C_1","C_2","C_3","C_4","C_5","C_6"), 
                               group2=c("D_1","D_2","D_3","D_4","D_5","D_6"), 
                               equal.disp = FALSE, smoothing = FALSE)
#Smoothing = FALSE for RRBS data due to non-continuous CpG site coverage.

#View top few results for each analysis:
head(DMLtest_female.exp)
head(DMLtest_male.exp)

##Sort the dieldrin-related DMC data frames by p-value (and remove NAs):
DMLtest_female.exp_sort1 <- DMLtest_female.exp[order(DMLtest_female.exp$pval, na.last=NA), ]
DMLtest_male.exp_sort1 <- DMLtest_male.exp[order(DMLtest_male.exp$pval, na.last=NA), ]

#View first few significant results ('top hits')
head(DMLtest_female.exp_sort1)
head(DMLtest_male.exp_sort1)

#Subset data frame based on FDR cutoff of 0.05
DMLtest_female.exp_sort2 <- subset(DMLtest_female.exp_sort1, fdr < 0.05)
DMLtest_male.exp_sort2 <- subset(DMLtest_male.exp_sort1, fdr < 0.05)
head(DMLtest_female.exp_sort2) #View first few significant results
head(DMLtest_male.exp_sort2)
summary(DMLtest_female.exp_sort2) #View summary of data frame variables
summary(DMLtest_male.exp_sort2)

#Institute more stringent cutoffs for DMC-calling -- at least 5% change in methylation by exposure cutoff
DMLtest_female.exp_sort3 <- subset(DMLtest_female.exp_sort2, abs(diff) > 0.05)
DMLtest_male.exp_sort3 <- subset(DMLtest_male.exp_sort2, abs(diff) > 0.05)
#Note: None of the DMCs with FDRs<0.05 were removed, indicating that all of these had an absolute difference of >5% methylation.

#View first few significant results
head(DMLtest_female.exp_sort3) 
head(DMLtest_male.exp_sort3)
```


## 3.DMR calling on exposure-related DMCs.
```{r DMR calling on called DMCs, echo=TRUE, warning=FALSE, message=FALSE, cache=FALSE}

#DMR calling using built in DSS 'callDMR()' function:
dmrs_female.exp1 <- callDMR(DMLtest_female.exp_sort1, p.threshold=0.01, minlen=20, minCG=3) #  6 DMRs
dmrs_female.exp2 <- callDMR(DMLtest_female.exp_sort2, p.threshold=0.01, minlen=20, minCG=3) # 1 DMRs
dmrs_female.exp3 <- callDMR(DMLtest_female.exp_sort3, p.threshold=0.01, minlen=20, minCG=3) # 1 DMRs
dmrs_male.exp1 <- callDMR(DMLtest_male.exp_sort1, p.threshold=0.01, minlen=20, minCG=3) # 4 DMRs
dmrs_male.exp2 <- callDMR(DMLtest_male.exp_sort2, p.threshold=0.01, minlen=20, minCG=3) # 0 DMRs
dmrs_male.exp3 <- callDMR(DMLtest_male.exp_sort3, p.threshold=0.01, minlen=20, minCG=3) # 0 DMRs
#Can manually adjust the p-value threshold for DMR calling.
dmrs_female.exp4 <- callDMR(DMLtest_female.exp_sort1, p.threshold=0.05, minlen=20, minCG=3) # 13 DMRs
dmrs_female.exp5 <- callDMR(DMLtest_female.exp_sort2, p.threshold=0.05, minlen=20, minCG=3) # 1 DMRs
dmrs_female.exp6 <- callDMR(DMLtest_female.exp_sort3, p.threshold=0.05, minlen=20, minCG=3) # 1 DMRs
dmrs_male.exp4 <- callDMR(DMLtest_male.exp_sort1, p.threshold=0.05, minlen=20, minCG=3) # 14 DMRs
dmrs_male.exp5 <- callDMR(DMLtest_male.exp_sort2, p.threshold=0.05, minlen=20, minCG=3) # 0 DMRs
dmrs_male.exp6 <- callDMR(DMLtest_male.exp_sort3, p.threshold=0.05, minlen=20, minCG=3) # 0 DMRs
#Can also manually adjust the length threshold for DMR calling.
dmrs_female.exp7 <- callDMR(DMLtest_female.exp_sort1, p.threshold=0.05, minlen=10, minCG=3) # 16 DMRs
dmrs_female.exp8 <- callDMR(DMLtest_female.exp_sort2, p.threshold=0.05, minlen=10, minCG=3) # 1 DMRs
dmrs_female.exp9 <- callDMR(DMLtest_female.exp_sort3, p.threshold=0.05, minlen=10, minCG=3) # 1 DMRs
dmrs_male.exp7 <- callDMR(DMLtest_male.exp_sort1, p.threshold=0.05, minlen=10, minCG=3) # 14 DMRs
dmrs_male.exp8 <- callDMR(DMLtest_male.exp_sort2, p.threshold=0.05, minlen=10, minCG=3) # 0 DMRs
dmrs_male.exp9 <- callDMR(DMLtest_male.exp_sort3, p.threshold=0.05, minlen=10, minCG=3) # 0 DMRs

#Notes: Different p-value thresholds were attempted. For final DMR calling, minimum length was set to 10 bp; minimum # of CpGs was set to 3.

#################################################################################################
#DMR calling from DSS gave only a few DMRs, so we also used an additional DMR calling function -- DMRcate:
source("https://bioconductor.org/biocLite.R")
biocLite("DMRcate")
library(DMRcate)

#First annotate the detected CpGs.
wgbsannot.female <- cpg.annotate("sequencing", DMLtest_female.exp)
wgbsannot.male <- cpg.annotate("sequencing", DMLtest_male.exp)

detectCores() #Use this to determine whether parallel processing is okay to attempt. If not, stick with mc.cores =1.

#Notes about DMRcate: The values of lambda and C should be chosen with care. For array data, the DMRcate authors currently #recommend that half a kilobase represents 1 standard deviation of support (lambda=1000 and C=2), and 20bp
#(C=50) for WGBS data. For RRBS data, we tried a variety of parameters, settling on the settings below:

#Final settings used to call DMRs using DMRcate.
wgbs.female.DMRs <- dmrcate(wgbsannot.female, lambda = 500, C = 4, min.cpgs = 3, mc.cores = 1)
wgbs.female.DMR.ranges <- extractRanges(wgbs.female.DMRs, genome = "mm10")
female_DMRs_DMRcate <- as.data.frame(wgbs.female.DMR.ranges) #9 DMRs w/ Stouffer < 0.1

setwd("/path/to/working/directory")
write.csv(female_DMRs_DMRcate, file = "DSS_DMRs_P3_Female_RRBS_Dieldrin_DMRcate_082418.csv")

wgbs.male.DMRs <- dmrcate(wgbsannot.male, lambda = 500, C = 4, min.cpgs = 3, mc.cores = 1)
wgbs.male.DMR.ranges <- extractRanges(wgbs.male.DMRs, genome = "mm10")
male_DMRs_DMRcate <- as.data.frame(wgbs.male.DMR.ranges) # 0 DMRs w/ Stouffer < 0.1

setwd("/path/to/working/directory")
write.csv(male_DMRs_DMRcate, file = "DSS_DMRs_P3_Male_RRBS_Dieldrin_DMRcate_082418.csv")

```

## 4. Write CSVs for results from models sorted by p-values.
```{r DMR calling on called DMCs, echo=TRUE, warning=FALSE, message=FALSE, cache=FALSE}
setwd("/path/to/working/directory")
write.csv(DMLtest_female.exp_sort3, file = "DSS_DMCs_P3_Female_RRBS_Dieldrin_081318.csv")
write.csv(dmrs_female.exp7, file = "DSS_DMRs_P3_Female_RRBS_Dieldrin_081318.csv")
write.csv(DMLtest_male.exp_sort3, file = "DSS_DMCs_P3_Male_RRBS_Dieldrin_081318.csv")
write.csv(dmrs_male.exp7, file = "DSS_DMRs_P3_Male_RRBS_Dieldrin_081318.csv")
```

## 5. Examine the test statistics and p-values for the female and male datasets.
```{r P values and test statistics, echo=TRUE, warning=FALSE, message=FALSE, cache=FALSE}
par(mfrow=c(1,2))
hist(DMLtest_female.exp_sort3$stat, 50, main="Combined Exposure test statistics", xlab="")
hist(DMLtest_female.exp_sort3$pval, 50, main="Combined Exposure P-values", xlab="")
hist(DMLtest_female.exp_sort3$fdr, 50, main="Combined Exposure FDRs", xlab="")

par(mfrow=c(1,2))
hist(DMLtest_male.exp_sort3$stat, 50, main="Combined Exposure test statistics", xlab="")
hist(DMLtest_male.exp_sort3$pval, 50, main="Combined Exposure P-values", xlab="")
hist(DMLtest_male.exp_sort3$fdr, 50, main="Combined Exposure FDRs", xlab="")
```

## 6. Annotate DMCs/DMRs using annotatr.
```{r P values and test statistics, echo=TRUE, warning=FALSE, message=FALSE, cache=FALSE}
###############BED format:
#In excel, remove top row of variable names and fix chromosome numbers, then save file as .txt file. File should be in standard BED3 format plus any additional columns of sample information.
#This file can then be read in by annotatr to annotate gene regions

##Annotate CpG sites using the annotatr package
source("https://bioconductor.org/biocLite.R")
biocLite("annotatr")
install.packages("stringi")
biocLite("regioneR")
biocLite("TxDb.Mmusculus.UCSC.mm10.knownGene")
biocLite("org.Mm.eg.db")
library(annotatr)
library(rtracklayer)
library(regioneR)
library(TxDb.Mmusculus.UCSC.mm10.knownGene)
library(org.Mm.eg.db)

################# Adding custom miRNA and enhancer region annotations to annotatr #################

library(GenomicRanges)
library(rtracklayer)
library(annotatr)

setwd("/path/to/working/directory")
mirna_gff <- import.gff("mmu_miRNA_annotations.gff3") # Imports mus musculus miRNA GFF3 genome annotation file (from: http://www.mirbase.org/ftp.shtml).
mirna_gff[1:4,]

#Coerce miRNA database to a data frame in R
mirna_gff_df <- as.data.frame(mirna_gff)

#Write .csv of data frame for conversion into BED6 format in excel:
setwd("/path/to/working_directory")
write.csv(mirna_gff_df, "miRNA_database_annotations_081418.csv")
#Convert .csv file to BED format in excel, and then save as .txt file.

## Custom annotation objects are given names of the form genome_custom_name
setwd("/path/to/working_directory")
mirna_db_file = "miRNA_database_annotations_081418_BEDformat.txt"
extraCols0 = c(gene_id='character', symbol='character', type='character')
#Use the read_annotations function to create custom annotation file for miRNA:
read_annotations(con = mirna_db_file, genome = 'mm10', name = 'miRNA',  extraCols = extraCols0, format='bed')

#Also create custom annotation for predicted enhancers identified from postnatal mouse midbrain (ENCODE).
#Raw file downloaded from: https://www.encodeproject.org/annotations/ENCSR114ZIJ/
setwd("/path/to/working_directory")
enhancer_db_file = "ENCFF129YCY_ENCODE_mouse_midbrain_enhancers_091318.bed"
extraCols0 = c(gene_id='character', symbol='character', type='character')
read_annotations(con = enhancer_db_file, genome = 'mm10', name = 'enhancer',  extraCols = extraCols0, format='bed')

#Double-check that custom annotation track has been successfully added to the mm10
print(annotatr_cache$get('mm10_custom_miRNA')) 
print(annotatr_cache$get('mm10_custom_enhancer')) 

##################### Annotating Data ##############################

#Note for annotatr: Used BED6+3 format file in the following format:
#chr4	  152512417	152512418	hyper	-0.953057058	0.033368133	0.986425191
#chr12	48140364	48140365	hyper	-0.932408194	0.044534154	0.976942348

#Columns 4, 5, 6, and 7 refer to direction of change, magnitude of change (diff),
#mu1, and mu2, respectively.

#Note that the start column is set to -1 compared to the end column; this 
#ensures that the final form of the data will have identical start and end values
#aftering reading in the data using annotatr.

#Set working directory and read in bed file
setwd("/path/to/working_directory")
dmc_female_file = "DSS_DMCs_P3_Female_RRBS_Dieldrin_081318_BEDformat.txt"
dmc_male_file = "DSS_DMCs_P3_Male_RRBS_Dieldrin_081318_BEDformat.txt"

#Define and rename extra columns in BED file:
extraCols = c(diff_meth = 'numeric', mu1 = 'numeric', mu2 = 'numeric')
dmc_female_regions = read_regions(con = dmc_female_file, extraCols = extraCols, genome = 'mm10',
                                  rename_name = 'DM_status', format = 'bed')
dmc_male_regions = read_regions(con = dmc_male_file, extraCols = extraCols, genome = 'mm10',
                                rename_name = 'DM_status', format = 'bed')
head(dmc_female_regions)
head(dmc_male_regions)

#Build annotations for mm10 based on variable of interest (dieldrin exposure):
builtin_annotations()
annots_exp = c('mm10_basicgenes','mm10_cpgs', 'mm10_lncrna_gencode', 'mm10_custom_miRNA', 'mm10_custom_enhancer')
annotations_exp = build_annotations(genome = 'mm10', annotations = annots_exp)

#Annotate the regions:
dmc_annots_female_exp = annotate_regions(
  regions = dmc_female_regions,
  annotations = annotations_exp,
  ignore.strand = TRUE)

dmc_annots_male_exp = annotate_regions(
  regions = dmc_male_regions,
  annotations = annotations_exp,
  ignore.strand = TRUE)

# Note: annotatr output is a special type of dataframe using the dplyr package.
# By default, dplyr::tbl_df objects have nice printing properties, but it
# hides extra columns that would ordinarily wrap. You can see them all with:
head(as.data.frame(dmc_annots_female_exp))
head(as.data.frame(dmc_annots_male_exp))

#Write the gene annotations to a tab-delimited text file:
setwd("/path/to/working/directory")
write.table(as.data.frame(dmc_annots_female_exp),file="DSS_DMCs_P3_RRBS_Dieldrin_Female_Annotated_091318.txt",sep="\t",quote=F)
write.table(as.data.frame(dmc_annots_male_exp),file="DSS_DMCs_P3_RRBS_Dieldrin_Male_Annotated_091318.txt",sep="\t",quote=F)

##Distributions of annotated DMCs

#Define order of differentially methylated CpGs (DMCs)
dm_order = c(
  'hyper',
  'hypo')
cpg_order = c(
  'mm10_cpg_inter',
  'mm10_cpg_islands',
  'mm10_cpg_shores',
  'mm10_cpg_shelves',
  'mm10_genes_promoters',
  'mm10_genes_3UTRs',
  'mm10_genes_5UTRs',
  'mm10_genes_exons',
  'mm10_genes_introns',
  'mm10_lncrna_gencode',
  'mm10_custom_enhancer') #NOTE: Tried including custom miRNA track, but none of CpGs annotated to any miRNAs, so I removed this track from the graphs (otherwise it shows as NA).
#NOTE: Tried including custom enhancer track from ENCODE. There are indeed CpG sites that map to enhancers!

#Plot categorical to view CpG distributions vs. random distribution.
dmc_female_exp = plot_categorical(
  annotated_regions = dmc_annots_female_exp,
  x = 'DM_status',
  fill = 'annot.type',
  x_order = dm_order,
  fill_order = cpg_order,
  position = 'fill',
  legend_title = 'knownGene Annotations',
  x_label = 'DM status',
  y_label = 'Proportion')
plot(dmc_female_exp)

#No 5'UTRs in male annotations, so the list needs to be adjusted:
dm_order1 = c(
  'hyper',
  'hypo')
cpg_order1 = c(
  'mm10_cpg_inter',
  'mm10_cpg_islands',
  'mm10_cpg_shores',
  'mm10_cpg_shelves',
  'mm10_genes_promoters',
  'mm10_genes_3UTRs',
  'mm10_genes_exons',
  'mm10_genes_introns',
  'mm10_lncrna_gencode',
  'mm10_custom_enhancer') #NOTE: Tried including custom miRNA track, but none of CpGs annotated to any miRNAs, so                          I removed this track from the graphs (it was showing as NA).
#NOTE: Tried including custom enhancer track from ENCODE. There are indeed some CpG sites that map to enhancers!

dmc_male_exp = plot_categorical(
  annotated_regions = dmc_annots_male_exp,
  x = 'DM_status',
  fill = 'annot.type',
  x_order = dm_order1,
  fill_order = cpg_order1,
  position = 'fill',
  legend_title = 'knownGene Annotations',
  x_label = 'DM status',
  y_label = 'Proportion')
plot(dmc_male_exp)

#Save Exposure-based DMC distribution R plot to ONES server:
setwd("/path/to/working/directory")
pdf("DSS_DMCs_P3_RRBS_Dieldrin_Female_Distribution_091318.pdf") 
plot(dmc_female_exp)
dev.off()

setwd("/path/to/working/directory")
pdf("DSS_DMCs_P3_RRBS_Dieldrin_Male_Distribution_091318.pdf") 
plot(dmc_male_exp)
dev.off()

#################### Repeat this process for the DMRs ####################

#Read in DMRs
setwd("/path/to/working/directory")

#DSS callDMR() DMRs
dmr_female_file = "DSS_DMRs_P3_Female_RRBS_Dieldrin_081318_BEDformat.txt"
dmr_male_file = "DSS_DMRs_P3_Male_RRBS_Dieldrin_081318_BEDformat.txt"

#Define and rename extra columns in BED file:
extraCols = c(diff_meth = 'numeric', mu1 = 'numeric', mu2 = 'numeric')
dmr_female_regions = read_regions(con = dmr_female_file, extraCols = extraCols, genome = 'mm10',
                                  rename_name = 'DM_status', format = 'bed')
dmr_male_regions = read_regions(con = dmr_male_file, extraCols = extraCols, genome = 'mm10',
                                rename_name = 'DM_status', format = 'bed')
head(dmr_female_regions)
head(dmr_male_regions)


#DSS + DMRcate DMRs:
setwd("/path/to/working/directory")
dmr_female_file2 = "DSS_DMRs_P3_Female_RRBS_Dieldrin_DMRcate_BEDformat_082418.txt"
dmr_male_file2 = "DSS_DMRs_P3_Male_RRBS_Dieldrin_DMRcate_BEDformat_082418.txt"
#DMRcate DMRs (BED3 format; No extra columns.)
dmr_female_regions2 = read_regions(con = dmr_female_file2, genome = 'mm10',
                                   format = 'bed')
dmr_male_regions2 = read_regions(con = dmr_male_file2, genome = 'mm10',
                                 format = 'bed')
head(dmr_female_regions2)
head(dmr_male_regions2)


#Build annotations for mm10 based on variable of interest (age):
annots_exp = c('mm10_basicgenes','mm10_cpgs', 'mm10_lncrna_gencode', 'mm10_custom_miRNA', 'mm10_custom_enhancer')
annotations_exp = build_annotations(genome = 'mm10', annotations = annots_exp)

##Now annotate!

#DSS DMRs:
dmrs_female_exp_annotated = annotate_regions(regions = dmr_female_regions, annotations = annotations_exp, ignore.strand = TRUE)

dmrs_male_exp_annotated = annotate_regions(regions = dmr_male_regions, annotations = annotations_exp, ignore.strand = TRUE)

#DMRcate DMRs:
dmrs_female_exp_annotated2 = annotate_regions(regions = dmr_female_regions2, annotations = annotations_exp, ignore.strand = TRUE)

dmrs_male_exp_annotated2 = annotate_regions(regions = dmr_male_regions2, annotations = annotations_exp, ignore.strand = TRUE)

# Note: annotatr output is a special type of dataframe using the dplyr package.
# By default, dplyr::tbl_df objects have nice printing properties, but it
# hides extra columns that would ordinarily wrap. You can see them all with:
head(as.data.frame(dmrs_female_exp_annotated))
head(as.data.frame(dmrs_male_exp_annotated))

## You could write the gene region annotations to a tab-delimited text file with:
write.table(as.data.frame(dmrs_female_exp_annotated),file="DSS_DMRs_P3_RRBS_Dieldrin_Female_Annotated_091318.txt",sep="\t",quote=F)

write.table(as.data.frame(dmrs_male_exp_annotated),file="DSS_DMRs_P3_RRBS_Dieldrin_Male_Annotated_091318.txt",sep="\t",quote=F)

#DMRcate DMRs:
setwd("/path/to/working/directory")
write.table(as.data.frame(dmrs_female_exp_annotated2),file="DSS_DMRs_P3_RRBS_Dieldrin_Female_DMRcate_Annotated_091318.txt",sep="\t",quote=F)
write.table(as.data.frame(dmrs_male_exp_annotated2),file="DSS_DMRs_P3_RRBS_Dieldrin_Male_DMRcate_Annotated_091318.txt",sep="\t",quote=F)

##Distributions of annotated DMRs were not visualized due to very low number.

```


## 7. Visualize identified DMRs.
```{r Data visualization, echo=TRUE, warning=FALSE, message=FALSE, cache=FALSE}

#First, need to install RnBeads (and dependencies):
source("https://bioconductor.org/biocLite.R")
biocLite("RnBeads")
biocLite("FDb.InfiniumMethylation.hg19")
biocLite("RnBeads.mm10")#Companion package for mouse mm10 genome:
biocLite("Gviz")
library(RnBeads)
library(RnBeads.mm10)
library(Gviz)

#RnBeads provides functionality for generating plots of methylation levels in custom genomic
#intervals similar to current genome browsers. It makes use of the ggbio package.

#Probably the most convenient and safe way to load bisulfite sequencing data is to use the 
#format presets which are tailored to particular tools and pipelines. 
#These presets are specified in the package option 'import.bed.style'. 

# Presets within RnBeads include bismark coverage files -- 'bismarkCov' 
#import.bed.style = 'bismarkCov'

######### Step 1 -- Data loading ############################

#The data.source argument of the rnb.run.import() and rnb.run.analysis() functions for
#analyzing bisulfite sequencing data requires a vector or list of length 1, 2 or 3 containing:

#1. the directory where the methylation bed files are located;
#2. the sample annotation sheet (based on RnBeads documentation -- http://rnbeads.mpi-inf.mpg.de/examples.php)
#3. the index of the column of the sample annotation sheet that contains the names or full
#paths to the bed files

library(RnBeads)
# Directory where your data is located
setwd("/path/to/working/directory")
getwd()
dataDir <- file.path(getwd())
sampleSheet <- file.path(dataDir, "sample_annotation_P3_RRBS_Female_Dieldrin_081418.csv") #Female samples with coverage files
sampleSheet2 <- file.path(dataDir, "sample_annotation_P3_RRBS_Male_Dieldrin_081418.csv") #Male samples with coverage files

######### Step 2 - Parameter settings for import ###################################

#Set global options prior to data import
rnb.options(
  identifiers.column = "sampleID",
  import.bed.style = "bismarkCov",
  assembly= "mm10"
)

#Enable large analysis of massive datasets on machines with intermediate RAM:
rnb.options(disk.dump.big.matrices=TRUE)
#The file system location which will be used by the ff objects is controlled by a global R option
#(notice that it is not an RnBeads option!):
options(fftempdir="/path/to/temp/directory/")
#By default, the ff files will be saved to the current R temporary directory, which is usually a
#subdirectory of the systems global temporary directory and can be found by executing:
tempdir()
#You can monitor the amount of hard drive space occupied by the RnBeads objects
#in the logger statements by setting:
rnb.options(logging.disk=TRUE)
#In order to ensure that the disk-backed objects are effectively cleaned up during the pipeline
#execution use the following option:
rnb.options(enforce.memory.management=TRUE)
#This will force RnBeads to use garbage collection so that no obsolete ff files remain on the
#hard drive.

################# Step 3 - Data import ############################
dataSource <- c(dataDir, sampleSheet) #Female P3 data
dataSource2 <- c(dataDir, sampleSheet2) #Male P3 data
rnbs <- rnb.execute.import(dataSource, data.type="bs.bed.dir") #rnbset created from P3 RRBS data
rnbs2 <- rnb.execute.import(dataSource2, data.type="bs.bed.dir") #rnbset created from BPA data

#Visualize beta value distribution for RRBS data:
rnb.plot.betadistribution.sampleGroups(meth(rnbs),rnb.sample.groups(rnbs)[[1]], "Exposure")
rnb.plot.betadistribution.sampleGroups(meth(rnbs2),rnb.sample.groups(rnbs2)[[1]], "Exposure")
#Beta distribution [0,1]. Beta-binomial model 
#Note: skewed towards low methylation; LOTS of beta values near zero!

################ Step 4 - Creating a genome browser figure ############

#Visualizing RRBS DMRs

#Start by visualizing Female RRBS data for DMR identified using DSS.
#Example: DMR annotated to the Grb10 gene prmoter/intron (ONLY IN MALES).

#First, visualize the female RRBS data at the Grb10 DMR identified from DSS:
chrom <- "chr11"
start <- 12026499	
end <- 12026597
sample.grouping <- rnb.sample.groups(rnbs)[[2]] #group by sample description -- corresponds to "exposure" (Control vs. Dieldrin)
#Adjust the color scheme of graph for easier interpretation:
rnb.options(colors.gradient = c("#ffffff", "#252525")) #eight class grey scale for methylation datan from ColorBrewer
rnb.options(colors.category = c("#b2182b", "#2166ac")) #two class blue gradient for category.
#Plot the graph
rnb.plot.locus.profile(rnbs, chrom, start, end, grps=sample.grouping, cvals.grps = rnb.getOption("colors.category"), cvals.meth = rnb.getOption("colors.gradient"))

#Now try visualizing the Male RRBS data for the Grb10 DMR identified using DSS
#As before (in the female data), DMR is annotated to the Grb10 gene promoter/intron.
chrom <- "chr11"
start <- 12026499	
end <- 12026597
sample.grouping <- rnb.sample.groups(rnbs2)[[2]] #group by sample description -- corresponds to "exposure" (Control vs. Dieldrin)
#Adjust the color scheme of graph for easier interpretation:
rnb.options(colors.gradient = c("#ffffff", "#252525")) #eight class grey scale for methylation datan from ColorBrewer
rnb.options(colors.category = c("#b2182b", "#2166ac")) #two class blue gradient for category.
#Plot the graph!
rnb.plot.locus.profile(rnbs2, chrom, start, end, grps=sample.grouping, cvals.grps = rnb.getOption("colors.category"), cvals.meth = rnb.getOption("colors.gradient"))

#To save PDF of DMR visualization:
pdf("P3_RRBS_Dieldrin_Male_Grb10_DMR_081418.pdf")
rnb.plot.locus.profile(rnbs2, chrom, start, end, grps=sample.grouping, cvals.grps = rnb.getOption("colors.category"), cvals.meth = rnb.getOption("colors.gradient"))
dev.off()

```
````
