## Supplementary material for "Developmental Dieldrin Exposure Alters DNA Methylation at Genes Related to Dopaminergic Neuron Development and Parkinson’s Disease in Mouse Midbrain": Supplemental Figures S1+S2+S3_Kochmanski_121118.docx

**Low-dose Developmental Dieldrin Exposure Alters Epigenome-wide DNA Methylation in the Mouse Midbrain**

**Supplementary Material**

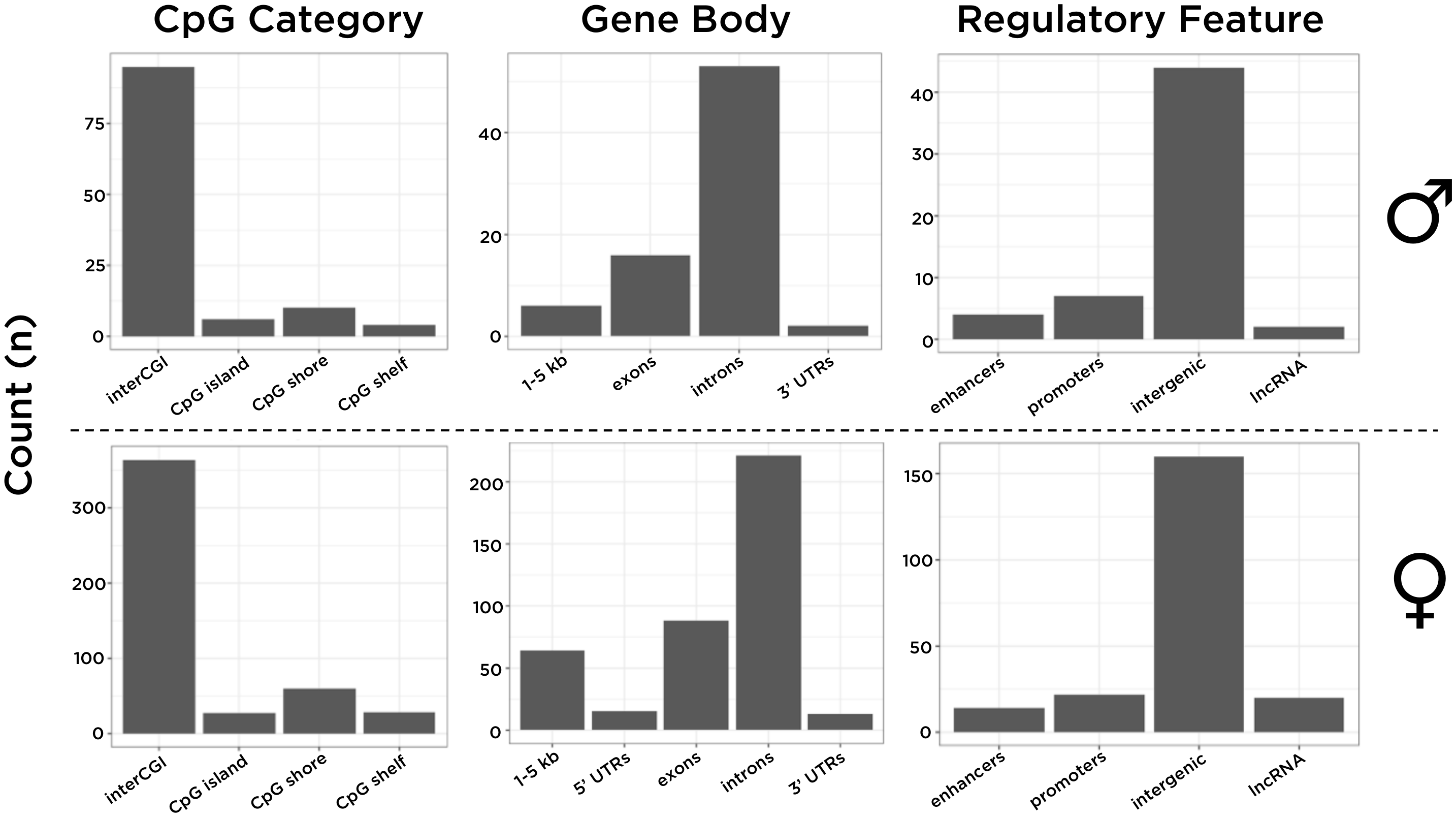

**Figure S1 – Distributions of male and female DMC annotations.** The *annotatr* R package was used to annotate DMCs and DMRs to CpG categories (e.g. CpG island), gene body locations (e.g. exon), and regulatory features (e.g. promoter). No DMCs annotated to 5’ UTRs in the male mice, so that category is not included in the male distribution. Both male and female DMCs showed similar genomic distributions of differential methylation, indicating that the types of genomic features affected by dieldrin exposure were not necessarily different by sex.

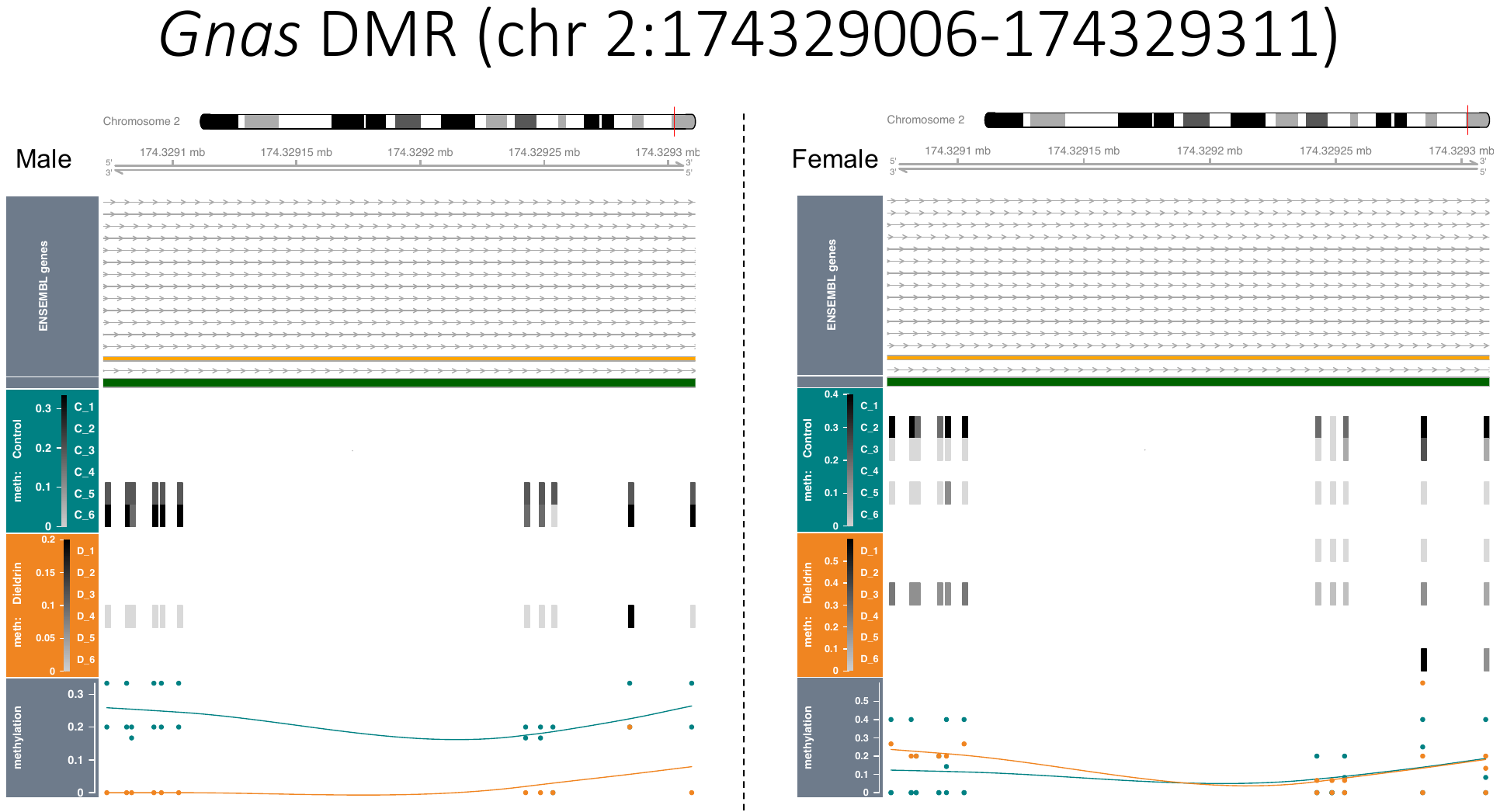

**Figure S2 – Differentially methylated region at the *Gnas* locus.** The male-specific DMR identified at the *Gnas* locus (chr 2:174329006-174329311) is driven by sparse data coverage across samples (n=3 and n=4 for male and female, respectively) and may be a spurious result. Shown here from top to bottom: Ensembl gene cartoons, CpG islands (green bars), and DNA methylation levels by individual sample in both a heat plot and dot plot with smoothed mean line. As indicated by sex labels, male data are shown on the left, while female data are shown on the right.

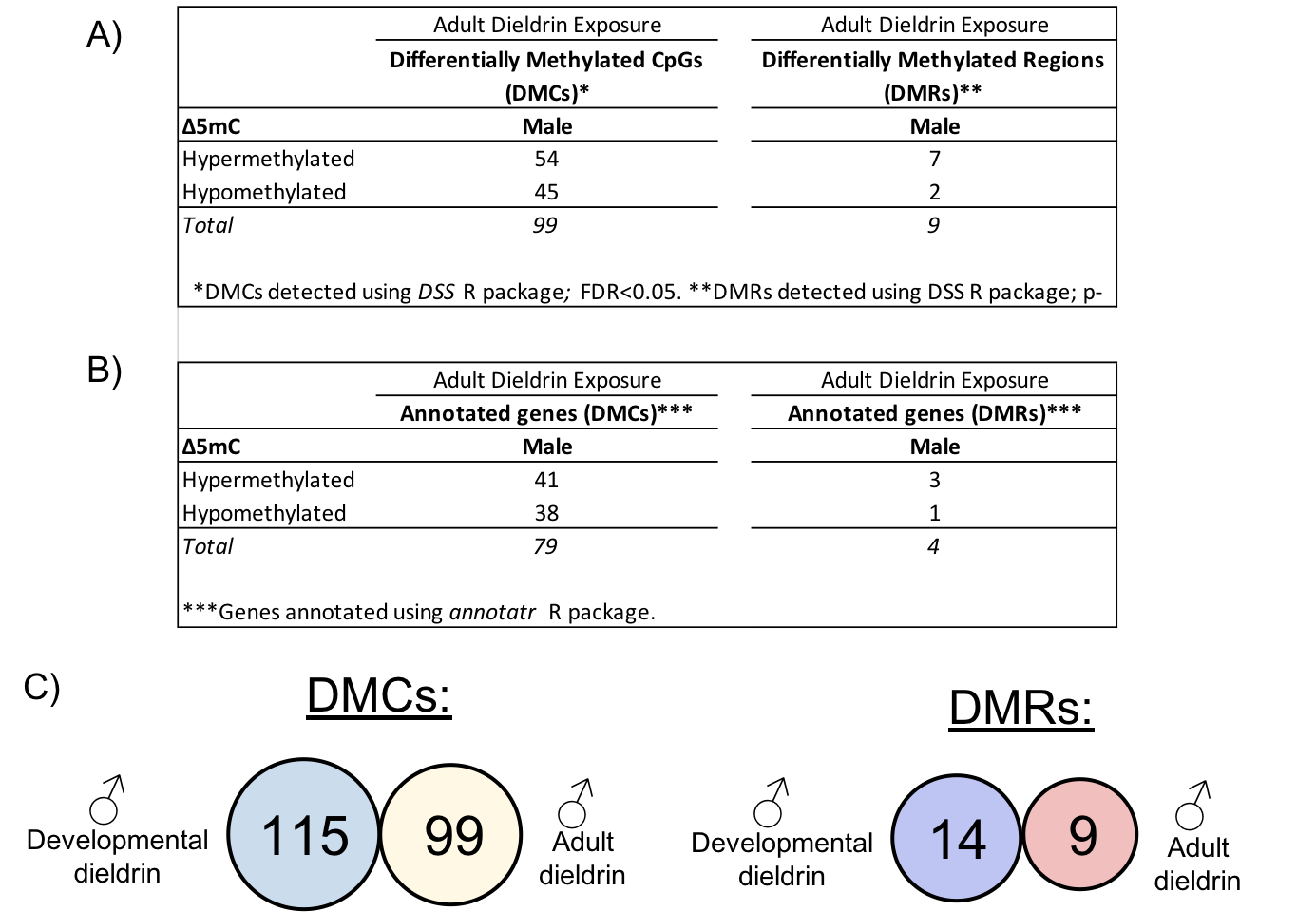

**Figure S3 - Differential methylation by adult dieldrin exposure.** Using the *DSS* R package, two-group Wald test models were used to test for differential DNA methylation by adult dieldrin exposure in RRBS data generated from male midbrain samples collected at 12 weeks of age. A) In the adult exposure study, we identified 99 differentially methylated CpGs, of which 54 were hypermethylated and 45 were hypomethylated. In addition to the DMCs, 9 DMRs were detected by adult dieldrin exposure. B) The identified DMCs and DMRs annotated to 79 and 4 unique genes, respectively. Some DMCs annotated to more than one gene, and 33 DMCs did not annotate to any known genes. C) None of the adult male exposure DMCs or DMRs overlapped with the developmental male exposure model results.
